# Bank vole immunoheterogeneity may limit Nephropatia Epidemica emergence in a French non-endemic region

**DOI:** 10.1101/130252

**Authors:** Adélaïde Dubois, Guillaume Castel, Séverine Murri, Coralie Pulido, Jean-Baptiste Pons, Laure Benoit, Anne Loiseau, Latifa Lakhdar, Maxime Galan, Philippe Marianneau, Nathalie Charbonnel

**Affiliations:** INRA, Centre de Biologie pour la Gestion des Populations, Montferrier sur Lez, France; ANSES, Unité de Virologie, Laboratoire de Lyon, France; ANSES, Plateforme d’Expérimentation Animale, Laboratoire de Lyon, France; CNRS, Laboratoire de Biométrie et Biologie Evolutive, Villeurbanne, France; CIRAD, Centre de Biologie pour la Gestion des Populations, Montferrier sur Lez, France

**Keywords:** *Myodes glareolus*, Rodent, Puumala hantavirus, Population genetics, Experimental infections, ecoimmunology

## Abstract

Ecoevolutionary processes affecting hosts, vectors and pathogens are important drivers of zoonotic disease emergence. In this study, we focused on nephropathia epidemica (NE), which is caused by Puumala hantavirus (PUUV) whose natural reservoir is the bank vole, *Myodes glareolus*. Despite the continuous distribution of the reservoir in Europe, PUUV occurence is highly fragmented. We questioned the possibility of NE emergence in a French region that is considered to be NE-free but that is adjacent to a NE-endemic region. We first confirmed the epidemiology of these two regions using serological and virological surveys. We used bank vole population genetics to demonstrate the absence of spatial barriers that could have limited dispersal, and consequently, the spread of PUUV into the NE-free region. We next tested whether regional immunoheterogeneity could impact PUUV chances to establish, circulate and persist in the NE-free region. Immune responsiveness was phenotyped both in the wild and during experimental infections, using serological, virological and immune related gene expression assays. We showed that bank voles from the NE-free region were sensitive to experimental PUUV infection. We observed high levels of immunoheterogeneity between individuals and also between regions. In natural populations, antiviral gene expression (*Tnf* and *Mx2* genes) reached higher levels in bank voles from the NE-free region. During experimental infections, anti-PUUV antibody production was higher in bank voles from the NE endemic region. Altogether, our results indicated a lower susceptibility to PUUV for bank voles from this NE-free region, what might limit PUUV circulation and persistence, and in turn, the risk of NE.

**Graphical abstract**

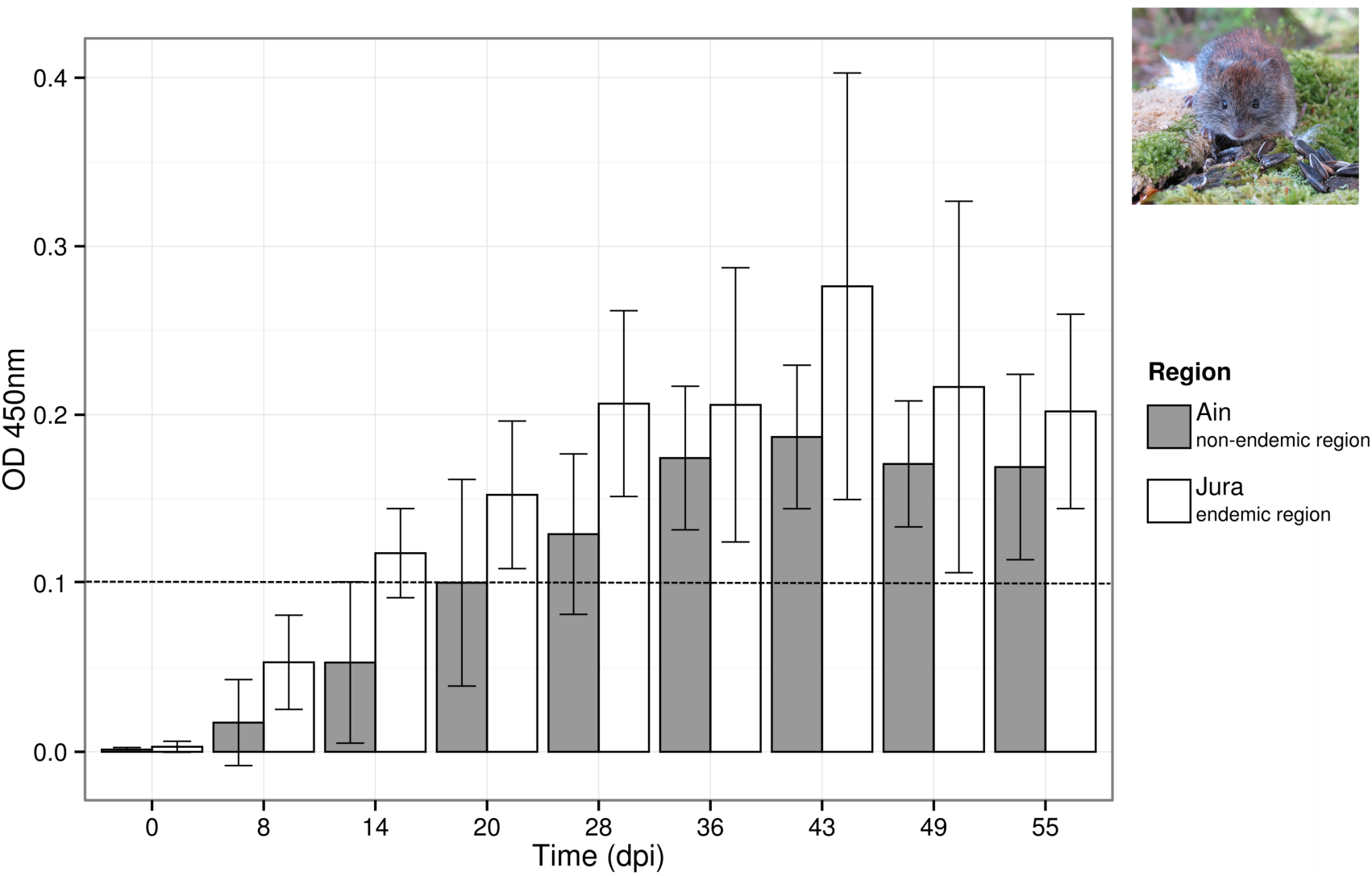

## 1. Introduction

Emerging infectious diseases (infections that have newly appeared in a population, or have existed but are rapidly increasing in incidence or geographic range, \(Morse and Schluederberg, 1990) have become an important public health threat these last decades (Daszak et al., 2000; King et al., 2006). The need for integrating evolutionary concepts to better understand and prevent them has recently been advocated (Vander Wal et al., 2014). Hence, hosts, vectors and pathogens evolve in response to environmental selective pressures but also as a result of inter-specific interactions, what, in turn, strongly impacts infectious disease dynamics and emergence.

Among important zoonotic (re-)emerging agents are hemorrhagic fever-causing hantaviruses of the Bunyaviridae family (Vapalahti et al., 2003). They are mainly rodent-borne (Henttonen et al., 2008) and are responsible for two kinds of human syndromes, hemorrhagic fever with renal syndrome (HFRS), mainly in Eurasia, and hantavirus cardiopulmonary syndrome (HCPS) in the Americas. For twenty years, since reliable tools for hantavirus infection diagnostics have become available, the (re-)emergence of zoonoses associated with hantaviruses has been recognized as a growing public health concern worldwide. In Europe, an increase in amplitude and frequency of epidemics due to hantaviruses has been observed these last 20 years (Heyman et al., 2011). Moreover, the geographic distribution of hantaviruses is expanding (e.g. for Seoul virus Jameson et al., 2013; Mace et al., 2013). Although this may be potentially and partly explained by increased clinician awareness and easier access to diagnostic tests, a significant rise in the real number of human cases is very likely (Heyman et al., 2012; Reynes et al., 2015). In France, Puumala virus (PUUV) is the main agent of hantavirus infections. It induces an attenuated form of HFRS called nephropathia epidemica (NE). PUUV is transmitted to humans via aerosols contaminated by excreta of its unique and specific natural reservoir host, the bank vole *Myodes glareolus*, which is a forest dwelling rodent species (Brummer-Korvenkotio et al., 1982). Although *M. glareolus* exhibits a spatially continuous distribution all over France, except on the Mediterranean coast, human cases are mostly being reported in the northeast part of the country (National Reference Center for Hantavirus http://www.pasteur.fr/fr/sante/centres-nationaux-reference/les-cnr/hantavirus, but see Reynes et al., 2015). Elsewhere, PUUV can be absent or the prevalence of PUUV in bank vole populations can reach high levels without any human case being reported yet (Castel et al., 2015). Niche modeling approaches based on climatic and environmental factors have been conducted to better predict the spatial distribution of NE disease (Zeimes et al., 2015). They highlighted geographic areas of high risk of NE emergence, although neither seropositive bank voles nor human cases had been detected yet. They also failed to detect geographic areas with high levels of PUUV prevalence in reservoir populations and no human case reported. These results suggest that the drivers of NE emergence are just beginning to be elucidated, and that other important factors than abiotic ones, including evolutionary processes and PUUV and/or host genetic characteristics, could be relevant. In particular, selective pressures affecting host-pathogen interactions may promote the evolution of different defense strategies in hosts, ranging from resistance, *i.e.* the ability to reduce pathogen burden to tolerance, *i.e.* the ability to limit the damage caused by a given parasite burden (Schneider and Ayres, 2008; Raberg et al., 2009). Previous field and laboratory studies have revealed some variability in the susceptibility of bank voles to PUUV infection, as reflected by the probability to be infected with PUUV (Olsson et al., 2002; Kallio et al., 2006; Deter et al., 2008b; Dubois et al., 2017a) and the pattern of PUUV excretion once bank voles are infected (Hardestam et al., 2008; Voutilainen et al., 2015). Moreover, Guivier et al. (2010; 2014) have shown significant differences in the level of immune related gene expression in bank voles from NE endemic and non-endemic regions at the European and regional scales. It was suggested that this immunoheterogeneity could reflect some balance of resistance / tolerance to PUUV. Indeed PUUV infection in bank vole is chronic and mainly asymptomatic (but see Tersago et al., 2012) and mounting immune responses is energetically costly and can induce immunopathologies (e.g. in human infections, Vaheri et al., 2013). Guivier et al. (2010; 2014) therefore proposed that bank voles from NE endemic areas would be more tolerant to the virus as a result of co-adaptation whereas those from NE non-endemic ones would be more resistant.

In this context, we proposed to investigate the hypothesis that immunoheterogeneity between bank vole populations could influence the geographic distribution range of PUUV, using two complementary approaches, natural population surveys and experimental infections. We focused on bank vole populations settled on both parts of the southern limit of PUUV distribution in France (Fig. 1), *i.e.* in the NE endemic area (human cases regularly reported, Jura, see National Reference Center for Hantavirus data) and in the NE non-endemic area (no human case ever reported yet, Ain). Our first objective was to discard the possibility of a spatial barrier that could prevent or limit the transmission of PUUV from the Jura to the Ain bank vole populations. Our second objective was to test the assumption of an important regional immunoheterogeneity between bank vole populations that could reflect different levels of sensitivity to PUUV infection. We predicted bank voles from Ain to be less sensitive to PUUV infection than those from Jura. Under this hypothesis, we expected bank voles from Ain to exhibit lower probability of infection and higher levels of immune responses to PUUV infection, hence reflecting their higher levels of resistance to this hantavirus.

**Fig. 1.**
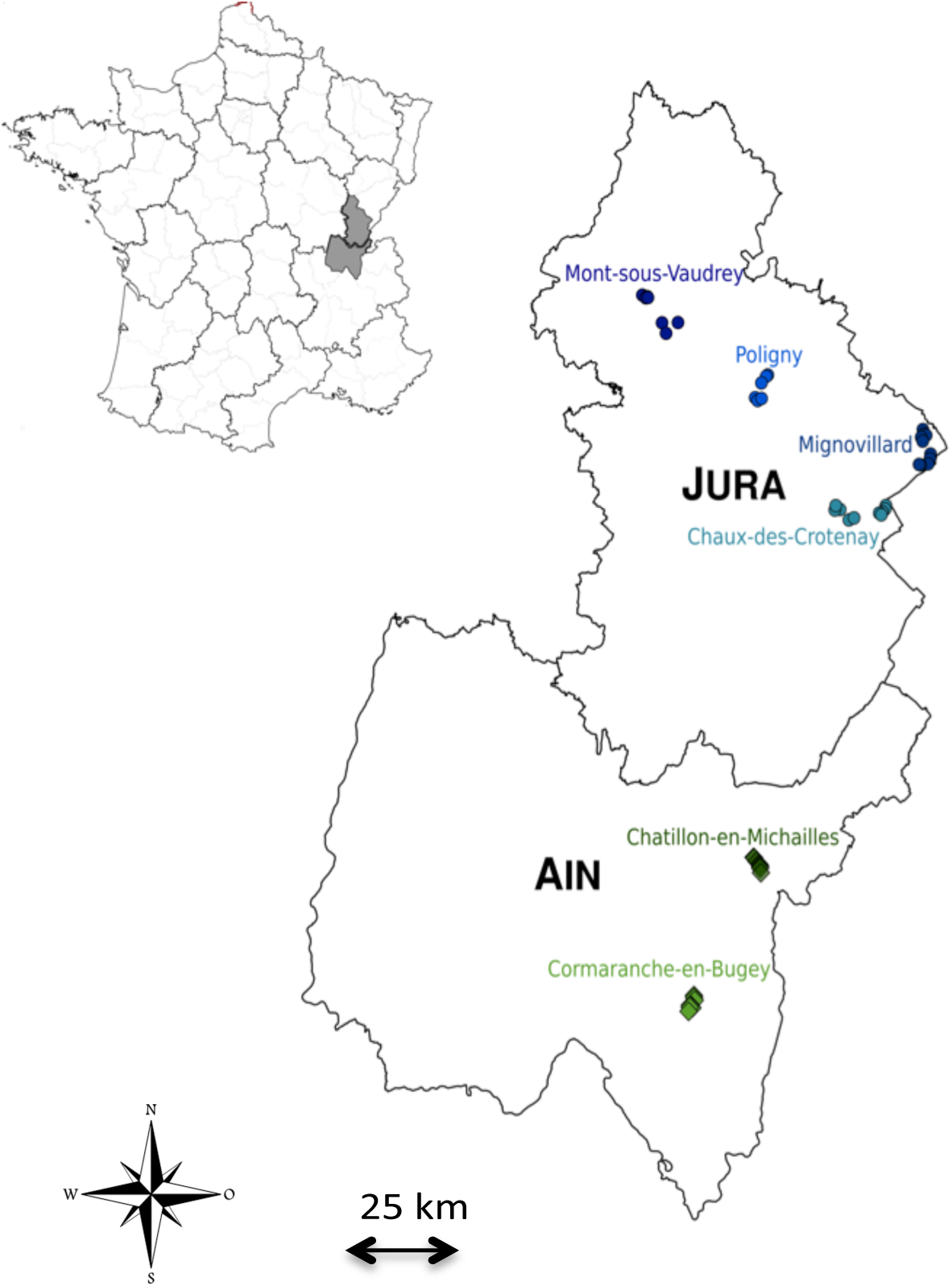
Map of the study area. Each color represents a site of trapping and each circle or diamond represents a trapping line. The location of the two regions (Jura and Ain) within France is shown on top left.

## 2. Material and methods

### 2.1. Ethical statement

All animal works have been conducted according to the French and European regulations on care and protection of laboratory animals (French Law 2001-486 issued on June 6, 2001 and Directive 2010/63/EU issued on September 22, 2010). Experimental protocols have been evaluated and approved by the Animal Ethics Committee C2EA-16 (ANSES/ENVA/UPEC, CNREEA n°16).

### 2.2 Natural population studies

#### 2.2.1. Rodent sampling

Bank voles were trapped in June and September 2014 in two adjacent regions of eastern France that have contrasted status in regard of human cases of NE. Jura is considered as an endemic zone as NE cases are regularly reported (National Reference Center for Hantavirus) whereas Ain is considered to be non-endemic as human case has never been reported yet (Fig. 1). These two areas are part of the same geological massif (Sommaruga, 1997) and they experience highly similar climatic conditions (WorldClim data, Supplementary Fig. 1).

Six sites were sampled in mixed forest of coniferous and deciduous tree species (Fig. 1, Table 1). The minimum distance between sites was 8.7 km. Six to ten lines of 20 live-traps (INRA) with about five meters interval were set up so that each sampling site consisted of a few km^2^ area. Traps were baited with sunflower seeds. Each trap was geolocated. The traps were checked daily and early in the morning. Trapping session per site lasted at least three nights so that a minimum of 35 voles were caught.

**Table 1.** Sampling site characteristics

Sampling site characteristics
| Region | Sites | Dates of sampling<br>(dd/mm/yyyy) | Number of<br>captured voles | Number of infected voles<br>(Number of infected<br>juveniles) | Prevalence<br>(Prevalence without<br>juveniles) |
| --- | --- | --- | --- | --- | --- |
| Ain | Chatillon-en-Michailles | 28/08/2014 - 31/08/2014 | 44 | 0 | 0 % |
| (non endemic) | Cormaranche-en-Bugey | 01/09/2014 - 03/09/2014 | 46 | 0 | 0 % |
| Jura<br>(endemic) | Chaux-des-Crotenay | 12/06/2014 - 15/06/2014<br>23/08/2014 | 56 | 4 (0) | 7.1 % |
|  | Mont-sous-Vaudrey | 07/06/2014 - 10/06/2014 | 40 | 4 (2) | 10.0 % (5 %) |
|  | Mignovillard | 16/06/2014 - 18/06/2014 | 45 | 0 | 0 % |
|  | Poligny | 24/08/2014 - 26/08/2014 | 46 | 4 (0) | 8.7 % |

Once trapped, animals were bled through retro-orbital sinus, killed by cervical dislocation, weighed, measured, sexed and dissected. Whole blood was stored at 4°C until centrifugation to collect sera 24 hours later. Sera, lungs and urine were immediately placed in dry ice and then stored at -80°C for virological and immunological analyses. Spleens were placed in RNAlater solution (Sigma) at 4°C during one night then stored at -20°C for population genetics and gene expression analyses.

#### 2.2.2. Microsatellite genotyping

Genomic DNA was extracted from a piece of spleen using the the EZ-10 96-Well Plate Genomic DNA Isolation Kit for Animal (BioBasic) according to manufacturer’s instructions, with a final elution of 400μL in elution buffer. Genotyping was performed at 19 unlinked microsatellites loci previously published by Rikalainen et al. (2008) using primers and cycling conditions described in Guivier et al. (2011) using an ABI3130 automated DNA sequencer (Applied Biosystems). Alleles were scored using GENEMAPPER software (Applied Biosystems).

#### 2.2.3. Serological and virological analyses

Serum samples were screened by IgG ELISA as described in Klingstrom et al. (2002). Due to the high cross-reactivity between the different hantavirus serotypes (Kruger et al., 2001), the plates were coated with Tula virus infected and non-infected cell lysates. Previous experiments have shown that anti-PUUV positive and negative sera reacted similarly with lysates of PUUV and TULV infected cells and with recombinant PUUV nucleocapsid protein (Dubois et al., 2017a).

Viral RNA was extracted from serum, lung and liver, which are target organs for PUUV (Gavrilovskaya et al., 1983; Bernshtein et al., 1999), and urine, as it is a main route of excretion for PUUV (Hardestam et al., 2008), using the QIAamp Viral Mini Kit (Qiagen). Quantitative RT-PCR (qRT-PCR) were performed using 2.5μL of viral RNA amplified using the SuperScript III One-Step RT-PCR system with Platinum Taq High Fidelity (Invitrogen) on LightCycler 480 (Roche Diagnostics) as described in Suppl. Table S1. Relative amounts of viral RNA (expressed in RNA copy per mg of tissue or per μl of liquid) were calculated using a standard curve obtained with *in vitro* transcribed RNA. All samples were tested in duplicate to avoid false positives and negatives.

#### 2.2.4. Immunological analyses

We quantified gene expression for three candidate immune related genes, namely *Tlr7*, *Tnf-α* and *Mx2*, that are relevant with regard to PUUV infections (Rohfritsch et al., 2013; Charbonnel et al., 2014; Dubois et al., 2017b). TLR7 is a receptor to virus and is probably involved in the detection of ssRNA viruses like hantaviruses (Bowie and Haga, 2005). The proinflammatory cytokine tumor necrosis factor alpha (TNF) and the antiviral protein Mx2 are known to limit PUUV replication in humans and in cell cultures (Kanerva et al., 1996; Temonen et al., 1996; Jin et al., 2001). We used *β-actin* as the endogenous reference gene, as previously validated by Friberg et al. (2011) in wood mice.

We analysed a subset of 170 bank voles including 10 adult PUUV-seropositive individuals and a number of PUUV-seronegative individuals that enabled to reach about 30 voles per site with an equal sex ratio. Total RNA was extracted from a piece of spleen using the NucleoSpin© 96 RNA kit (Macherey-Nagel) following manufacturer’s instructions. As secondary lymphoid organ, the spleen has important immune functions and is the site of low levels of PUUV replication (Korva et al., 2009). RNA extractions were electrophoresed on 1.5% agarose gels and visualized using ethidium bromide staining to check for quality, then total RNA concentrations were measured using a NanoDrop 8000 spectrophotometers (Thermo Scientific) and normalized to 200 ng/μL using RNase free water. For *β-actin, Tnf-α* and *Mx2*, we used specific primers described in previous studies (Guivier et al., 2010; Guivier et al., 2014). For *Tlr7*, we designed the following specific primers based on the cDNA consensus sequences obtained for 12 *M. glareolus* samples (GenBank Accession Numbers: KX463605 - KX463616): Tlr7-Mg2-F (5’-TACCAGGACAGCCAGTTCTA-3’), Tlr7-Mg2-R (5’-GCCTCTGATGGGACAGATA-3’). We generated cDNA from 4 μL of extracted RNA (800 ng per reaction), in a 20 μl reaction, using the Improm-II Reverse Transcription System (Promega), according to the conditions specified by the manufacturer for oligo (dT)15 primers. We performed a quantitative PCR on a LightCycler 480 (Roche Diagnostics), using the 384-multiwell plate format, as previously described in Guivier et al. (2010; 2014). PCR efficiencies were estimated using the LinRegPCR software (Ruijter et al., 2009). Candidate gene mRNA relative expression levels were estimated for each sample using the method developed by Pfaffl(2001).

### 2.3. Experimental studies

#### 2.3.1. Rodent sampling

Bank voles were trapped in May 2015 in two sites already sampled in 2014 (Cormaranche forest in Ain and Arbois forest in Jura) and using the same sampling protocol as described above. They were brought back to the laboratory, tested for the presence of anti-PUUV IgG, held during three weeks in quarantine and then retested for the presence of anti-PUUV IgG. Ten voles from Ain and nine voles from Jura, all of them being seronegative, were transferred in a ABSL-3 capacity and held individually in ISOcages N (Tecniplast). Food and water were provided *ad libitum*, with fresh vegetables once a week.

#### 2.3.2. Experimental infections

1.7x10^3^ f.f.u of PUUV Sotkamo strain 1:10 diluted in DMEM (ThermoFisher Scientific) were injected by subcutaneous route in each of the 19 bank voles. From 7 to 55 days post-infection (*dpi*), blood, saliva and feces were collected once a week for each bank vole. We did not succeed collecting urine samples. Approximately 200μL of whole blood was sampled through the retro-orbital sinus and stored at 4°C until centrifugation to collect sera 24 hours later. Saliva was collected using sterile swabs subsequently placed in 300μL of Hank’s Balanced Salt Solution (Life Technologies) and vortexed for 10 seconds. Feces were sampled directly from the anus. All samples were stored at -80°C until analyses. At the end of the experiment (55 *dpi*), all the bank voles were euthanized by cervical dislocation. Blood, lungs, liver, kidneys, urine and feces were collected and stored at -80°C until analyzes. Three bank voles, two from Ain and one from Jura, died during the course of experimental infection (at 28 *dpi*) and were not considered for further analyzes.

#### 2.3.3. Serological and virological analyses

PUUV IgG detection was performed on sera as described in 2.2.2. Viral RNA was extracted from all samples (sera, lung, liver, kidney, feces, saliva and urine) and PUUV RNA detection was realized using qRT-PCR for all the samples as described earlier. To improve the sensitivity of viral detection, nested RT-PCR were applied to sera, feces and saliva, using the Titan One Tube RT-PCR System (Roche) and Taq DNA Polymerase (Qiagen).

Gene expression could not be performed on bank voles from this experiment as spleens could only be collected at 55 *dpi.* It was too far from the infection to expect any difference in gene expression between bank voles (Hardestam et al., 2008; Schountz et al., 2012), and it did not enable to study gene expression kinetics.

### 2.4. Statistical analyses

#### 2.4.1. Natural population analyses

We analysed the genetic structure of bank vole populations using microsatellite data. We tested the conformity to Hardy-Weinberg equilibrium (HWE) for each locus and each population. We analyzed linkage disequilibrium (LD) for each pair of loci using Genepop v4.2 (Raymond and Rousset, 1995). Corrections for multiple tests were performed using the false discovery rate (FDR) approach as described in Benjamini and Hochberg (1995). We described the spatial genetic structure by estimating the genetic differentiation between each pair of sites using Weir and Cockerham’s pairwise *F*_*ST*_ estimates (1984). Significance was assessed using exact tests and FDR corrections. Several complementary analyses were performed to test for the existence of spatial barriers that could limit vole dispersal in this area, especially between Jura and Ain. We first computed a discriminant analysis of principal components (DAPC), which is a multivariate, model-free approach to clustering (Jombart et al., 2010). Because DAPC is sensitive to the number of principal components used in analysis, we used the function *optim.a.score* to select the correct number of principal components. We next performed an analysis of molecular variance (AMOVA) in Arlequin v3.5.1.2 (Excoffier and Lischer, 2010). Compared to the DAPC, it enabled to test specifically for genetic differences between regions, considering the potential variability between sites within each region. Finally models of isolation by distance (IBD) were applied to test for geographic-genetic correlations. IBD models were analyzed independently for each region and for the whole dataset, so that disruption of gene flow between regions could also be detected. Genetic differentiation was estimated for each pair of individuals and Mantel tests were applied to test for a correlation between matrices of genetic differentiation and of Euclidean distances between individuals, using 10000 permutations, in Genepop. We also calculated confidence intervals for the slope of the regression line by bootstrapping over loci (ABC intervals, Di Ciccio and Efron, 1996).

We tested for variations of serological and immunological characteristics in natural populations between regions using generalized linear mixed models with the GLMER function implemented in the LME4 package for R 3.1.0 (R Core Team, 2013). The fixed variables included in the models were *region, sex* and *weight* and all pairwise interactions. When they were non-significant, they were removed from the model. The factor *site* was included as a random factor. Chi-square tests with Bonferroni correction were applied to analyze the effect of significant variables using the package PHIA for R. We analysed independently PUUV serological status (using a binomial response variable) and each gene expression level *Qn* (using a log-transformed response variable for the three genes). To prevent confounding effects due to the potential induction of the three candidate genes expression following PUUV infection, we limited our statistical analyses to PUUV-seronegative bank voles (Guivier et al., 2014).

#### 2.4.2. Experimentation analyses

We tested for regional differences in bank vole responses to PUUV experimental infection by examining the levels of anti-PUUV IgG (*OD*_*450nm*_, using a normal response variable) between Jura and Ain through time. Generalized mixed linear models were applied as described above. The fixed variables included in the models were *region, dpi* and their interaction. Bank vole identity was included as a random effect. Possible variation in the level of viral RNA between regions was tested using a linear model with the fixed effect *region*.

## 3. Results

### 3.1. Natural population studies

#### 3.1.1. Population genetic structure

Three loci, Cg16H5, Cg16E4 and Cg6G11, were excluded from the genetic analyses due to the poor quality profiles obtained. Six bank voles were also excluded because they could not be amplified for most loci. Two other microsatellite loci, Cg2F2 and Cg3F12, showed deviations of HWE in four and six sites respectively, suggesting the presence of null alleles. They were removed from further analyses. Five out of 84 tests for deviation from HWE were significant with a different loci involved each time. 25 of 546 pairs of loci (4.58%) exhibited significant linkage disequilibrium but the loci involved were not consistent among sites. Thus, our final dataset included 271 bank voles genotyped at 14 microsatellites loci.

Spatial pairwise *F*_*ST*_ ranged between 0.009 and 0.03. All of these estimates were significant after FDR correction (Table 2). DAPC clustering showed that the genetic variation of the six sites overlapped (Fig. 2), although the first discriminant component tended to slightly separate the two regions. The second discriminant component revealed that one site in Jura (Mignovillard) tended to be different from the three others and that the two sites in Ain tended to be separated from each other (Fig. 2). AMOVA analyses revealed an absence of significant differentiation between the two regions, and significant variation between sites within regions (Table 3). IBD patterns were highly significant whatever the dataset considered. Genetic differentiation between bank voles decreased with spatial distance, and the slope of these regressions were similar, respectively 0.003, 0.004 and 0.005 for Ain, Jura and the whole dataset, confirming an absence of spatial barrier between the two regions (Table 4).

**Fig. 2.**
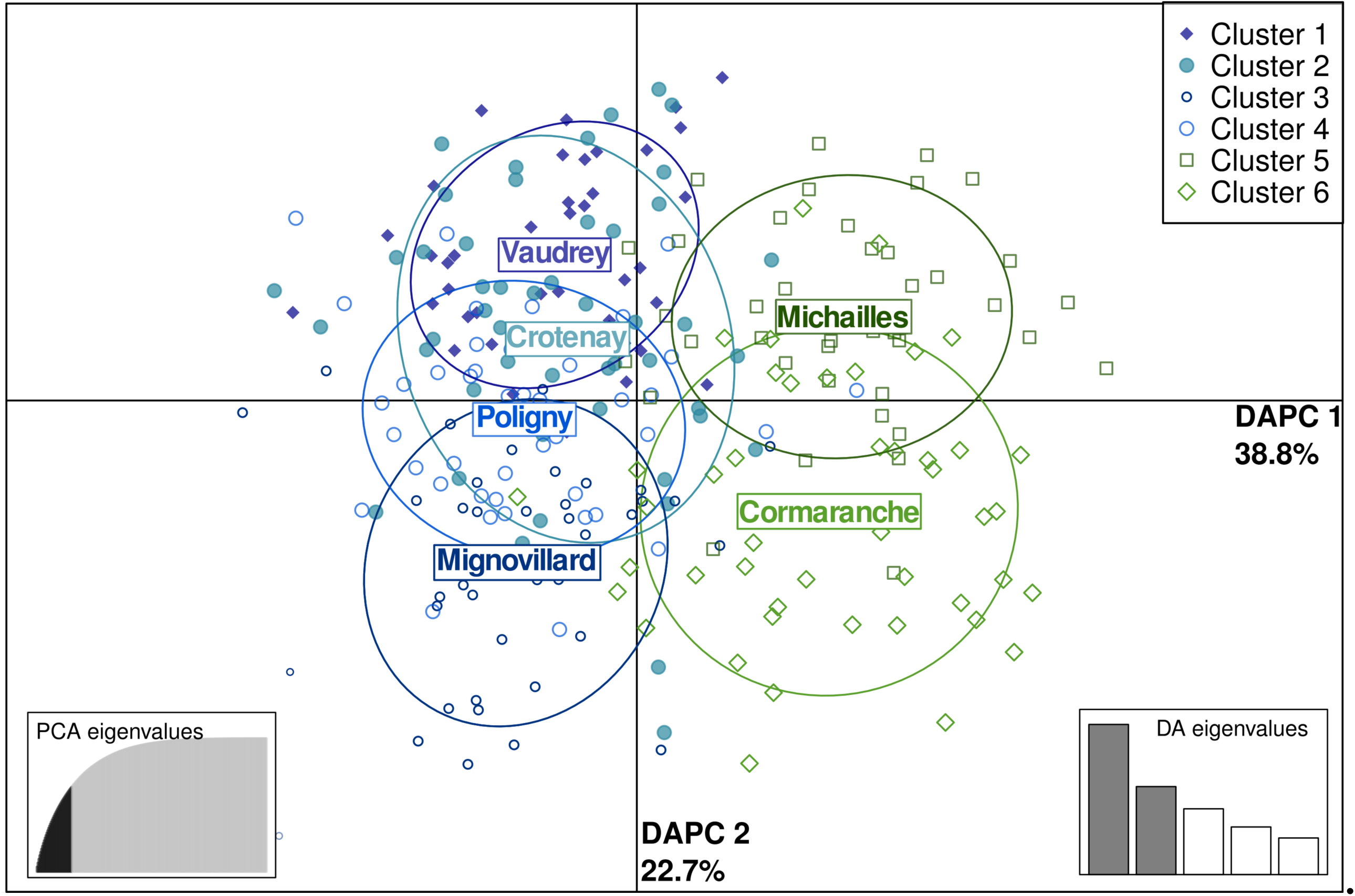
Scatterplot of the first two principal components discriminating *M. glareolus* populations by sites using a discriminant analysis of principal components (DAPC) based on microsatellite data. Points represent individual observations. Envelops represent population membership. Bank vole populations from Ain (NE non-endemic region) are represented in green and those from Jura (NE endemic region) in blue.

**Table 2.** *F*_*st*_ estimates between sites per year (lower side) and associated *p*-values of Fisher exact tests (upper side).

|  | Vaudrey | Crotenay | Mignovillard | Poligny | Chatillon | Cormaranche |
| --- | --- | --- | --- | --- | --- | --- |
| Vaudrey | | $< 10^{-5}$ | $< 10^{-5}$ | $< 10^{-5}$ | $< 10^{-5}$ | $< 10^{-5}$ |
| Crotenay | **0.01676\*\*\*** | | $< 10^{-5}$ | $< 10^{-5}$ | $< 10^{-5}$ | $< 10^{-5}$ |
| Mignovillard | **0.02296\*\*\*** | **0.01887\*\*\*** | | $< 10^{-5}$ | $< 10^{-5}$ | $< 10^{-5}$ |
| Poligny | **0.01501\*\*\*** | **0.00926\*\*\*** | **0.01453\*\*\*** | | $< 10^{-5}$ | $< 10^{-5}$ |
| Chatillon | **0.02142\*\*\*** | **0.01844\*\*\*** | **0.02279\*\*\*** | **0.02071\*\*\*** | | $< 10^{-5}$ |
| Cormaranche | **0.03116\*\*\*** | **0.02501\*\*\*** | **0.02161\*\*\*** | **0.02388\*\*\*** | **0.01566\*\*\*** |  |

Significant $p$ -values are in bold; \*\*\* $p \leq 0.001$

**Table 3.** Results of the analysis of molecular variance (AMOVA). The AMOVA hierarchy was based on geographical features, comparing the two regions (Ain and Jura) and the sites within the regions.

| Source of variation | *d.f.* | Variance of components | Percentage of variation | *p* |
| --- | --- | --- | --- | --- |
| Among regions | 1 | 0.04481 | 0.73 | 0.068 |
| Among sites within regions | 4 | 0.09903 | 1.61 | **< 10<sup>-5</sup> \*\*\*** |
| Within sites | 536 | 6.00519 | 97.66 | **< 10<sup>-5</sup> \*\*\*** |

Significant *p*-values are in bold; \*\*\* $p \leq 0.001$

**Table 4.** Isolation by distance characteristics — probability (IBD *p*), intercept and slope calculated for each region and for the whole dataset. 95% confidence intervals (CI) for the slope of the IBD were obtained by bootstrapping over the 14 microsatellite loci.

| Dataset | IBD $p$ | Intercept | Slope [95% CI] |
| --- | --- | --- | --- |
| Ain | **$< 10^{-5}$ \*\*\*** | 0.043 | 0.003 [$2 \cdot 10^{-5}$, 0.006] |
| Jura | **$< 10^{-5}$ \*\*\*** | 0.059 | 0.004 [$6 \cdot 10^{-4}$, 0.009] |
| Ain and Jura | **$< 10^{-5}$ \*\*\*** | 0.057 | 0.005 [0.003, 0.009] |

4 Significant $p$ -values are in bold; \*\*\* $p \leq 0.001$

#### 3.1.2. Variations of serological, virological and immunological features in natural populations

277 bank voles were trapped in the six sites surveyed (Table 1). Among them, 12 captured in Jura were PUUV seropositive. Two of these individuals had very low OD. Moreover, they had a low body mass, so that we thought they were juveniles carrying maternal antibodies and not PUUV infected individuals (weight lower than 17.5g, see Voutilainen et al., 2016). They were therefore excluded from further analyses. None of the bank vole captured in Ain was PUUV seropositive. Seroprevalence levels in Jura sites ranged between 7.1 % and 10.0 %, except in Mignovillard where no PUUV seropositive bank vole was detected. The best model that explained bank vole serological status only included *weight* (*X*^*2*^_*1*_ = 5.541, *p* = 0.019). Heavier bank voles were more likely to be PUUV seropositive.

The serum, lung, liver and urine of the 10 adult seropositive individuals were tested using qRT-PCR. Three sera were PUUV negative and the seven other samples showed viral RNA ranging between 1.01e4 and 1.58e5 copies.mg^-1^. The viral RNA load in the lung and liver ranged respectively between 7.50e6 and 4.88e9 copies.mg^-1^, and 1.79e6 and 8.63e8 copies.mg^-1^ (Suppl. Table S2). Urine samples could be collected for only five of these 10 adult bank voles. Two of these excreta were PUUV positive as their quantitative RT-PCR cycle thresholds, *C*_*T*_, were lower than 35 cycles. The viral load ranged between 2.08e4 and 2.26e4 copies.mg^-1^.

Extreme values of gene expression levels were detected for the three candidate genes (60% higher than the other values). The models were run with or without these extreme values. *Mx2* gene model revealed a significant difference between regions (without outliers: *X*^*2*^_*1*_ = 4.56, *p* = 0.033; with outliers: *X*^*2*^_*1*_ = 6.00, *p* = 0.01 Fig. 3a). Bank voles from Ain over-expressed *Mx2* compared to those from Jura. *Tlr7* gene model included the variable *sex* (without outliers: *X*^*2*^_*1*_ = 13.67, *p* = 2.10^-4^; with outliers *X*^*2*^_*1*_ = 7.80, *p* = 0.005, Fig. 3b), with higher values of *Tlr7* expression detected in females than in males. When including outliers, *Tnf-α* gene model revealed a significant effect of *region*, *weight* (*X*^*2*^_*1*_ = 9.173, *p* = 0.002; *X*^*2*^_*1*_ = 4.474, *p* = 0.034; respectively, Figs. 3 c, d) and of the interaction *sex * region* (*X*^*2*^_*1*_ = 5.783, *p* = 0.016). *Tnf-α* expression was higher in bank voles from Ain than those from Jura. This effect was stronger when considering males only. These results have to be taken cautiously as they turned to be non-significant when the outliers were not included in the model. Then we only found a significant effect of *weight* (*X*^*2*^_*1*_ = 11.50, *p* = 6.10^-4^), with heavier bank voles exhibiting higher levels of *Tnf-α* expression than lighter ones.

**Fig. 3.**
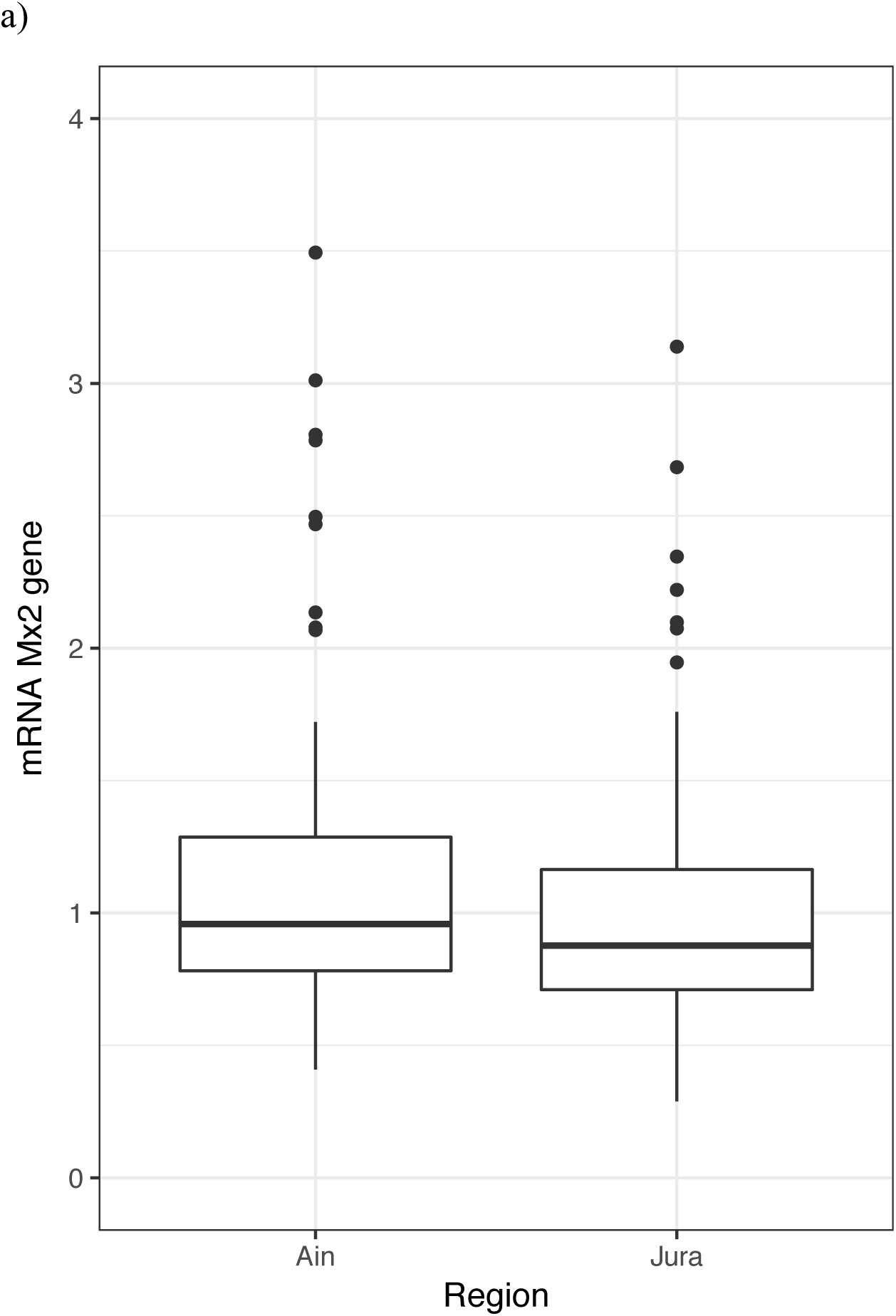
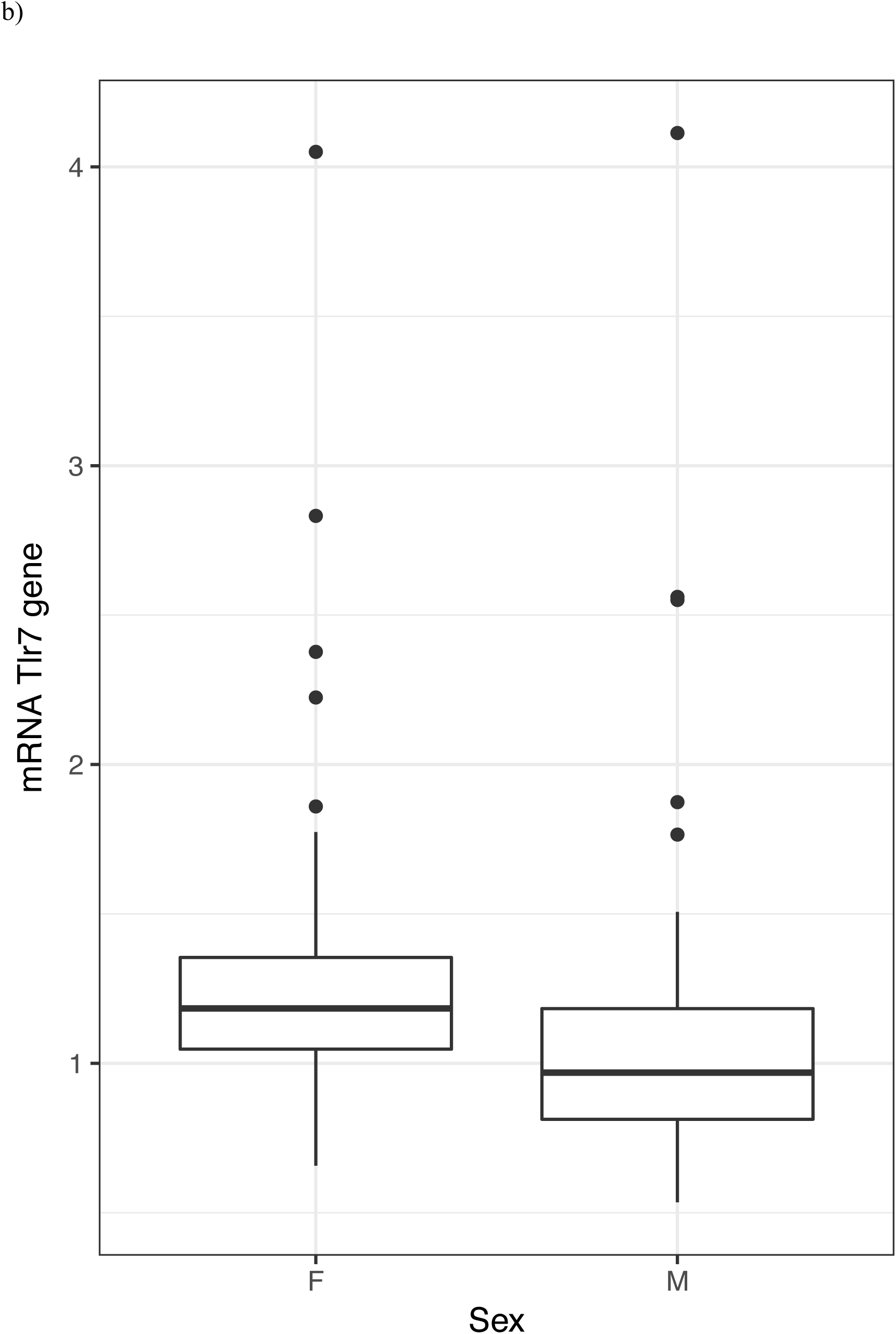
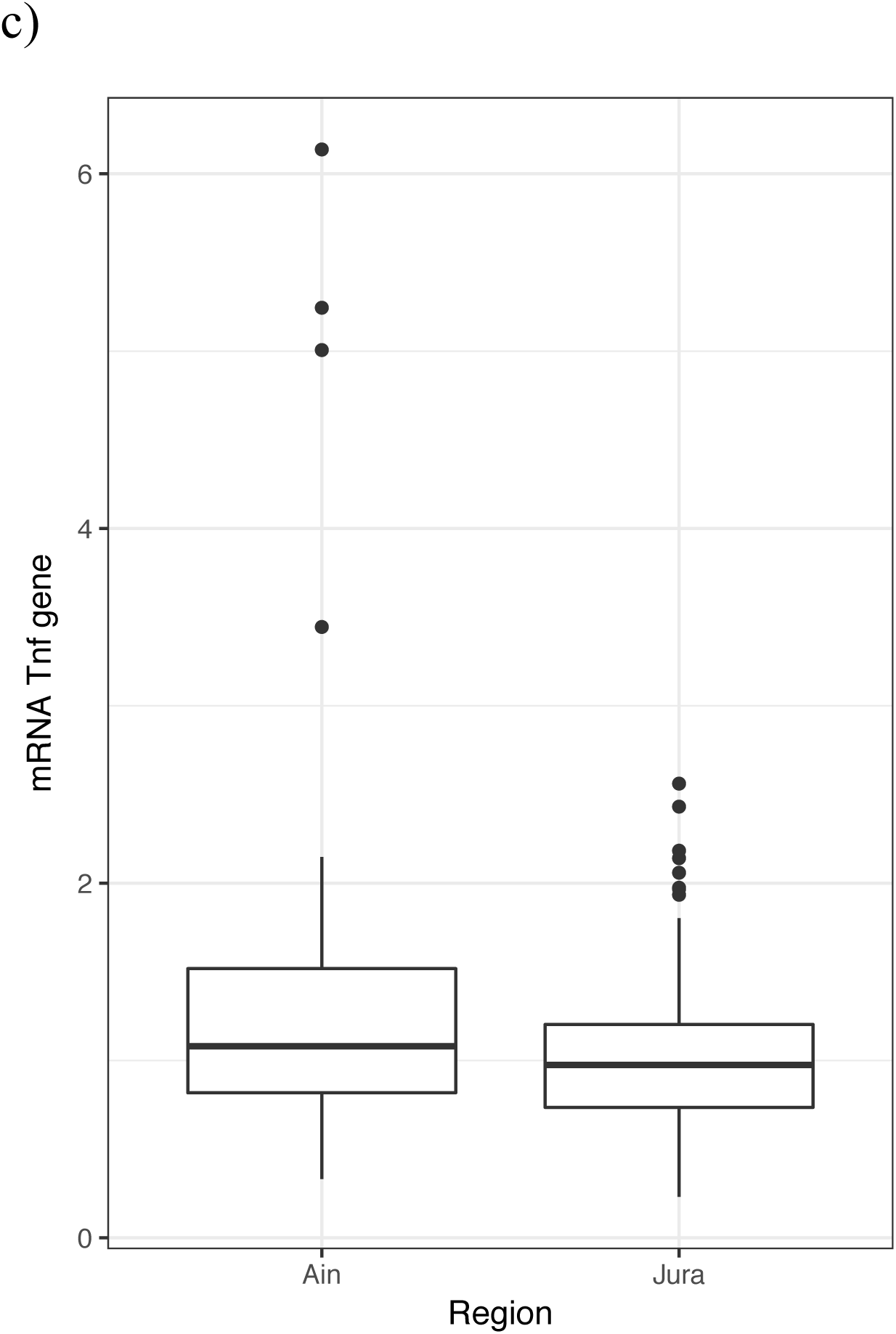
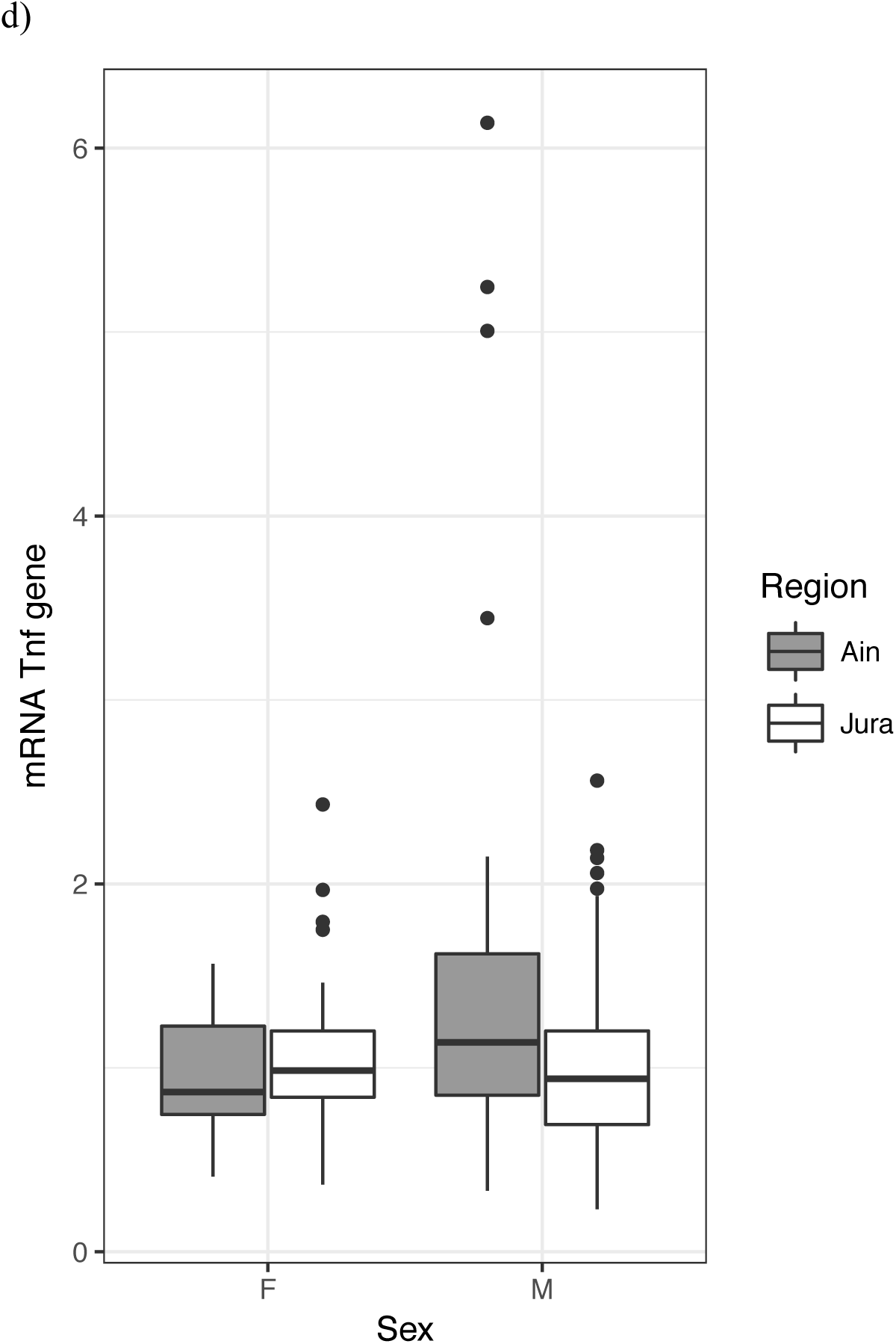
Variations of immunological characteristics in natural bank vole populations. a) Variations of *Mx2* gene expression between individuals of the two regions, Ain (NE non-endemic region) and Jura (NE endemic region). b) Variations of *Tlr7* gene expression between males and females. c) and d) Variations of *Tnf-α* gene expression between the two regions and between sex within the two regions.

### 3.2 Variations of serological and virological features during experimental infections

Three bank voles died before the end of the experiment and were not considered in the analyses. Among the 16 remaining bank voles experimentally infected with PUUV (eight from Ain and 8 from Jura), 14 seroconverted by the end of the study. The two bank voles that did not seroconvert at the end of the experiment (55 *dpi*) originated from Ain (Suppl. Table S3). Considering bank voles that seroconverted, our model showed a lower *OD*_*450nm*_ in voles from Ain than those from Jura (*X*^*2*^_*1*_= 4.126, *p* = 0.042, see Fig. 4), and a significant variation of *OD*_*450nm*_ through time (*X*^*2*^_*1*_ = 195.129, *p* = 2.10^-16^).

**Fig. 4.**
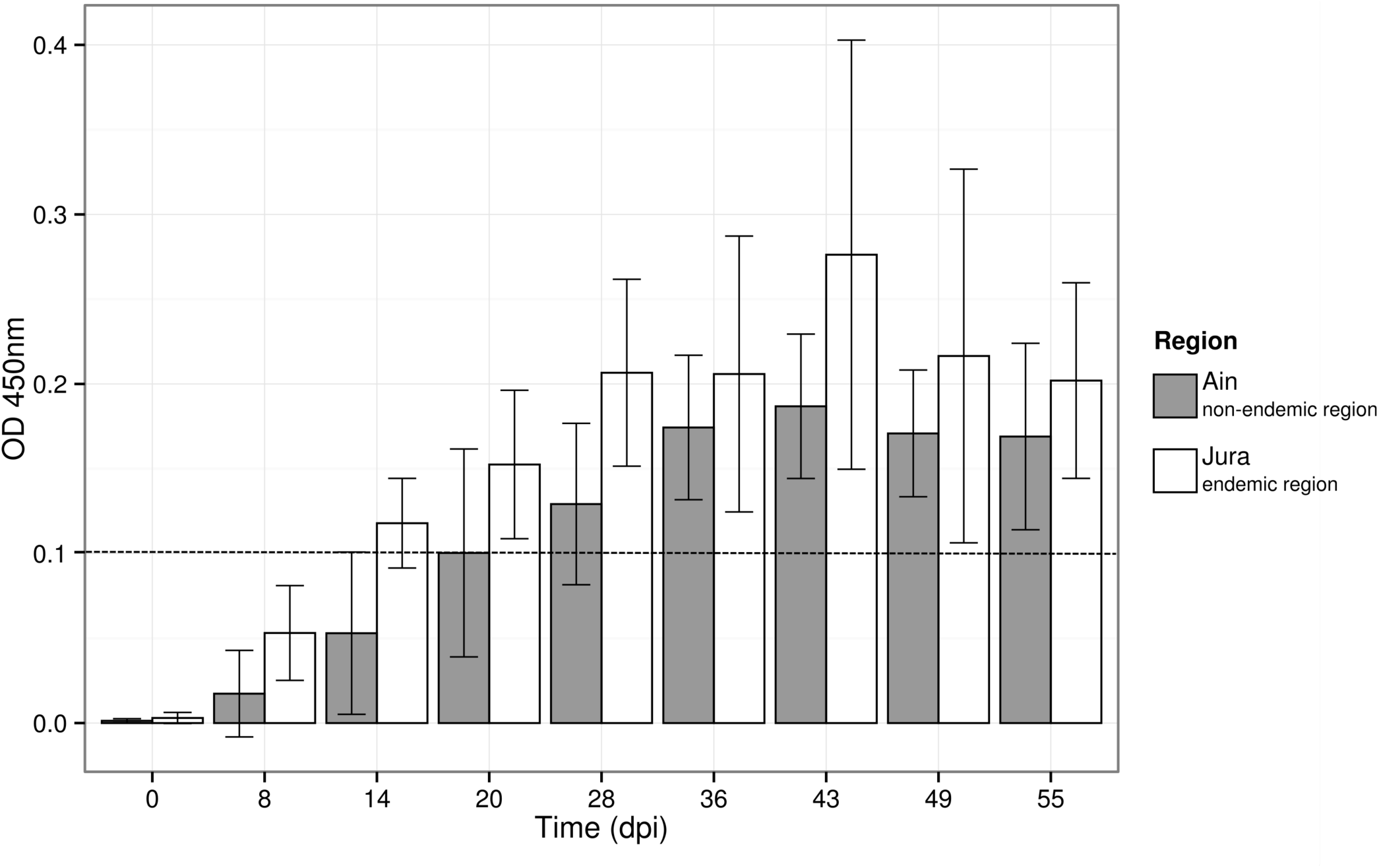
Variation of the optical density in ELISA test (*OD*_*450nm*_) over time for the 14 bank voles that seroconverted during the experimental infections. Barplots and error bars represent the mean *OD*_*450nm*_ value ± SD for all the individuals in each region. The hatched line represents the threshold above which the bank voles were considered to be PUUV seropositive.

Viral RNA was detected by qRT-PCR in the sera of a single bank vole from Ain at 14 and 20 *dpi*. Nested RT-PCR allowed detecting PUUV RNA at 8 *dpi* in the sera of two bank voles from Ain and of four bank voles from Jura (Suppl. Table S4). In addition, PUUV RNA could not be detected in any feces or saliva samples tested using qRT-PCR or nested RT-PCR.

At the end of the experiment (55 *dpi*), viral RNA was detected by qRT-PCR in the lungs of 13 bank voles (including one that did not seroconvert) over the 16 infected during the experiment. The relative amount of PUUV RNA ranged between 4.34e5 and 4.07e7 RNA copies. mg^-1^ (Suppl. Table S5). The statistical model revealed that PUUV viral load in lungs did not vary between the two regions. The amounts of viral RNA detected in the other organs were lower and concerned less individuals, in particular in liver: only four individuals showed positive results, and less than 3.07e6 RNA copies. mg^-1^ were detected (Suppl. Table S5). Finally, no viral RNA could be detected in saliva or urine samples collected at 55 *dpi*, and only two bank voles (one from each region) had PUUV RNA in their feces (Suppl. Table S5).

Finally, our results revealed a strong inter-individual variability in the serological / virological relationships. All bank voles but one that seroconverted had at least one organ/excreta with viral RNA at 55 *dpi*. Moreover, one bank vole that had not seroconverted at 55 *dpi* exhibited viral RNA in its lungs and kidneys (Suppl. Table S5).

## 4. Discussion

### 4.1 – Contrasted PUUV epidemiological situations between the French regions Ain and Jura

To our knowledge, this is the first bank vole serological study ever conducted in the French Ain region with regard to PUUV. It is probably because no NE case had been reported there yet (National Reference Center for Hantaviruses) and PUUV studies remained focused on endemic areas. Deepening our understanding of infectious disease emergence requires to broaden our research out of the areas where diseases are well described, to better identify the factors limiting or facilitating emergence success (e.g. for plague in Madagascar Gascuel et al., 2013; Tollenaere et al., 2013). None of the 90 bank voles captured in Ain during this field survey was seropositive for PUUV antibodies. This result tends to confirm the absence of PUUV circulation in these bank vole populations, or at such a low level that it can not be detected using ecological surveys. An extensive survey in more sites or in different years could still be performed to assert more firmly the absence of PUUV circulation in bank vole populations settled south from Jura. In the opposite, and as expected from the existence of known NE cases, we found PUUV seropositive bank voles in all but one site from Jura. Seroprevalence levels were very similar to those observed in the Ardennes epidemic region or in previous studies conducted in Jura (ranging from 0 to 18%, Guivier et al., 2011; Deter et al., 2008a). Although no bank vole from Mignovillard forest had anti-PUUV antibodies, we know from previous studies that PUUV may circulate in this site (Deter et al., 2008a). The existence of these two adjacent regions with contrasted epidemiological situations with regard to PUUV reinforced the need to better assess the risk of PUUV emergence and persistence in Ain. It emphasized the necessity to question whether PUUV infected bank voles from Jura could migrate to Ain, thus enhancing the chance for PUUV to be introduced there, and whether PUUV could persist in bank vole populations from Ain once introduced.

### 4.2 – High bank vole gene flow between the two French regions Ain and Jura

We developed a population genetic approach to evaluate the possibility that a spatial barrier could prevent bank vole gene flow from Jura to Ain, what would limit the chance of PUUV introduction in this latter region through the dispersal of infected animals. Indeed PUUV is transmitted directly between bank voles, from contaminated excreta or through bites, without any vector (Korpela and Lähdevirta, 1978; Yanagihara et al., 1985). PUUV introduction from one site to another may therefore strongly result from the introduction of an infected bank vole, either through wood transport or natural dispersal. For example, Guivier et al. (2011) showed that despite frequent local extinction of PUUV in hedge networks, the virus was frequently re-introduced in these habitats through the emigration of seropositive bank voles coming from close forests.

Our results revealed weak spatial genetic structure between sites and regions, with levels of genetic differentiation ranging between 1 and 3%. These estimates are very similar to the levels of genetic differentiation observed in previous studies conducted at the same geographical scale on bank voles using the same genetic markers (Guivier et al., 2011) or on other rodent species (Arvicola scherman, Berthier et al., 2005). These results revealed high population size and / or high gene flow between populations (Wright, 1951). We did not find any signature of disrupted gene flow between bank voles from the two regions. The spatial structure observed was mainly due to isolation by distance, *i.e.* migration rates being inversely proportional to the geographic distance between populations. This pattern is also frequently observed in micromammals, including rodents, when studied at this geographical scale (Aars et al., 2006; Berthier et al., 2006; Bryja et al., 2007). The fact that the rates of isolation by distance did not differ within region and over the whole dataset corroborated the hypothesis of an absence of spatial barrier disrupting gene flow between Jura and Ain. As such, the possibility for PUUV infected bank voles to disperse from Jura to Ain, and consequently, the possibility for PUUV introduction in Ain seemed likely.

Considering this possibility, the reason why PUUV, which is known to circulate in Jura since the early 2000’s, has not reached the Ain NE-free region needs to be questioned. It could be explained by the low progression of PUUV from northern to southern populations, and the emergence of NE in Ain would only be a question of time. It could also be due to differences in metapopulation functioning between bank vole populations in Ain and Jura. Higher genetic drift, lower migration rates between populations could account for lower probability of PUUV persistence in Ain (Guivier et al., 2011). However, we did not find any evidence supporting this hypothesis based on the population genetic analyses conducted using microsatellites. We therefore explored the possibility that bank voles from Ain and Jura could present different levels of sensitivity to PUUV infections, that would in turn lead to contrasted PUUV epidemiology in these two regions.

### 4.3 – High levels of immunoheterogeneity between bank voles

The assessment of immunological variations in wild populations has recently been at the core of eco-epidemiological studies, enabling the analysis of their environmental and evolutionary causes as well as the prediction of their epidemiological consequences (Jackson et al., 2011). Here we described strong individual variability of bank vole immune responses from both natural population survey and experimental infections. In the wild, susceptibility, defined here as the probability of being infected with PUUV, strongly depended on the weight of bank voles, which is a proxy for age and/or body condition. Heavier/older bank voles are likely to disperse more than younger ones and have accumulated more opportunities to be exposed to PUUV (see for details Voutilainen et al., 2016). In addition, these heavier/older bank voles could also suffer from immunosuppression due to high levels of corticosterone associated with breeding or to decreased body condition (Beldomenico et al., 2008). Moreover, we revealed variable amounts of viral RNA between bank voles, for all organs and excretas considered, what may result from differences in the time since infection or individual characteristics. The low number of animals that could be handled in highly secured animal facilities (ABSL3) did not enable to test the impact of individual factors on the probability of getting infected and on their specific sensitivity. Nevertheless, the experiments conducted allowed controlling for PUUV strain or quantity injected, and some variability was still detected both in anti-PUUV antibody production (*e.g.* 2 out of 16 bank voles did not seroconvert 55 days post-infection) and in the amount of viral RNA detected in organs and excretas. It was therefore relevant to analyse whether immune responsiveness also differed between bank voles. Considering three immune related candidate genes, namely *Tlr7, Mx2* and *Tnf-α*, we found that sex was an important factor mediating inter-individual immunoheterogeneity. In the wild, females over-expressed *Tlr7* gene compared to males. These observations corroborated common patterns observed in vertebrates showing that females are more immunocompetent than males (Klein and Flanagan, 2016). Indeed, TLR7 is involved in the recognition of diverse viruses (Heil et al., 2004; Lund et al., 2004) and impaired expression and signaling by TLR7 may contribute to reduced innate immune responses during chronic viral infections (Hirsch et al., 2010). The sexual dimorphism in *Tlr7* gene expression is observed in bank voles for the first time. This gene is linked to the X chromosome, and it belongs to the 15% of X-linked genes escaping inactivation and being expressed from both the active and inactive X chromosome (Plath et al., 2002). This could explain the over-expression of *Tlr7* gene observed in females compared to males. Surprisingly, we did not find any evidence of higher levels of *Tnf-α* gene expression in female compared to male bank voles, as was previously detected by Guivier et al. (2014). Higher expression levels were even detected in males compared to females in the region Ain. This pattern was driven by three bank voles with extreme values, that might correspond to recently infected males and blur the general picture observed in the Ardennes (Guivier et al., 2014). Altogether, this high inter-individual immunoheterogeneity, which was observed in the wild and confirmed when performing experimental infections, is important with regard to PUUV epidemiology as it highlighted bank voles with increased likelihood of being infected, excreting and further transmitting the virus (e.g. super-spreaders, Lloyd-Smith et al., 2005).

### 4.4. Regional bank vole immunoheterogeneity - an explanation for the contrasted PUUV epidemiology observed in adjacent French regions ?

Evidence of immunoheterogeneity was found between Jura and Ain, two regions with contrasted NE epidemiological statuses. It concerned several immune pathways, including the antiviral and adaptative immune responses of bank voles to PUUV infections. Bank voles from Ain mounted lower levels of anti-PUUV antibodies than those from Jura during PUUV experimental infections. Unfortunately, this regional pattern could hardly be interpreted in terms of rodent sensitivity to PUUV as the role of IgG during hantavirus infections remains unclear (review in Schonrich et al., 2008). Indeed, the presence of IgG can protect rodents from subsequent challenges but does not participate in eliminating the virus (review in Easterbrook and Klein, 2008; Schountz and Prescott, 2014). The similar amounts of viral RNA observed in bank voles from Ain and Jura at the end of the experiment could support the hypothesis that sensitivity to PUUV did not differ between these two regions. However, this result has to be taken cautiously as it is likely that variability in virus replication and excretion might only be visible sooner after the infection (Hardestam et al., 2008; Dubois et al., 2017a). Because we observed more bank voles from Jura than Ain with PUUV RNA in their sera eight days after the infection, we can not refute the possibility of a regional heterogeneity in sensitivity to PUUV.

The immune-related gene expression analyses conducted using wild bank voles also provide arguments in favor of this hypothesis. Rodents from Ain over-expressed *Mx2* and *Tnf-α* genes compared to bank voles from Jura. These results are congruent with previous studies conducted in the French Ardennes and over Europe (Guivier et al., 2010; Guivier et al., 2014). These immune related genes were chosen to represent some of the main pathways underlying bank vole immune responses to PUUV infections. They encode respectively for TNF, a proinflammatory cytokine and Mx2, an antiviral protein, that both limit PUUV replication (Kanerva et al., 1996; Temonen et al., 1996; Jin et al., 2001). Besides, the overproduction of these molecules could induce immunopathologies (Li and Youssoufian, 1997; Wenzel et al., 2005; Bradley, 2008), what could drive a balance of resistance / tolerance to PUUV (Guivier et al., 2010; Guivier et al., 2014). Altogether these regional differences in bank vole responses to PUUV could reflect potential divergence with respect to resistance and tolerance strategies. These differences may be mediated by a subset of individuals within populations, as suggested by bank voles exhibiting extreme values of immune related gene expression and mostly explaining the significant differences observed. Within population evolution from resistance to tolerance has previously been demonstrated in an unmanaged population of Soay sheep (Hayward et al., 2014) and in wild populations of field voles (Jackson et al., 2014). Further assessment of bank vole health and fitness would be required to confirm this hypothesis of tolerant and resistant phenotypes with regard to PUUV infections (Raberg et al., 2009). It would also be interesting to address the issue of potential physiological trade-offs between inflammatory and antibody-mediated responses (Lee and Klasing, 2004). The patterns observed in this study could suggest that bank voles from the NE endemic region would promote an adaptative immunity in response to PUUV infection instead of a more costly and potentially damaging inflammatory response, potentially as a result of co-adaptation between bank vole and PUUV (Easterbrook and Klein, 2008).

## Conclusions

This study highlighted the importance of combining natural population surveys and experimental approaches in addressing questions related to immunoheterogeneity and its potential consequences for epidemiological questions. Here the natural population survey enabled to study a large number of animals and to describe inter-individual and inter-population variability in immune responsiveness, that may further blur the results obtained from experimental infections. Besides, the experiments enabled to control for the time since exposure to PUUV and to follow immune responses kinetics. The diverse array of eco-evolutionary approaches developed here brought important answers to evaluate the global risk of NE emergence in a non endemic region. Our results supported the possibility for PUUV introduction in the region Ain via the dispersal of PUUV seropositive bank voles from Jura. They also indicated the possibility of PUUV circulation in this non endemic region as experimental infections revealed that bank voles from Ain are sensitive to PUUV. Therefore, NE emergence in this region might only be a question of time. But several arguments also indicated that PUUV persistence might be unlikely. Lower susceptibility to PUUV could account for the absence or low number of PUUV infected bank voles in the Ain region, and also potentially for the weak level of PUUV excretion in the environment. Further investigations are now required to identify the mechanisms underlying bank vole immunoheterogeneity between adjacent NE endemic and non endemic regions, in particular with regard to abiotic (e.g. climate, resource availability) or biotic conditions (e.g. spatially varying pathogen communities, see for the bank vole Behnke et al., 2001; Ribas Salvador et al., 2011; Razzauti-Feliu et al., 2015; Loxton et al., 2016). Sociological and human behavioral factors would then be the ultimate step to analyse what limits PUUV transmission between *M. glareolus* and humans.

## Acknowledgements

Data used in this work were partly produced through the genotyping and sequencing facilities of ISEM (Institut des Sciences de l’Evolution-Montpellier) and Labex CeMEB (Centre Méditerranéen Environnement Biodiversité). We are grateful to EU grants GOCE-CT-2003- 010284 EDEN and FP7-261504 EDENext, that have partially funded this research. A. Dubois PhD was funded by an INRA-EFPA / ANSES fellowship.

---

**Supplementary Table 1.**
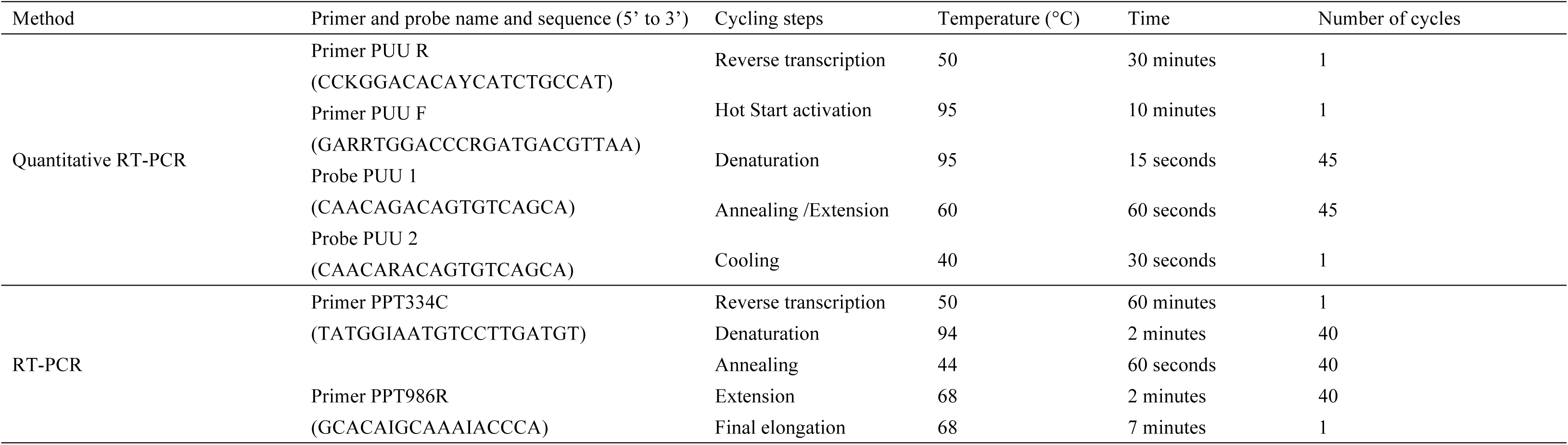
Sequences of primers and probes and cycling conditions of quantitative and nested RT-PCR.

**Supplementary Table 2.** Detection of PUUV RNA in sera, lung, liver and urinea of the 10 adult seropositive bank voles detected in natural populations.

| Region | Individual number | Sera (RNAcopies. $\mu\text{l}^{-1}$) | Lung (RNA copies. $\text{mg}^{-1}$) | Liver (RNA copies. $\text{mg}^{-1}$) | Urine (RNA copies. $\mu\text{l}^{-1}$) |
| --- | --- | --- | --- | --- | --- |
| Jura | NCHA00014 | N | 1.21e8 | 3.42e7 | N |
|  | NCHA00041 | N | 7.27e8 | 1.95e7 | N |
|  | NCHA00071 | 8.90e4 | 7.13e8 | 5.79e8 | N |
|  | NCHA00073 | 1.91e4 | 6.61e8 | 8.09e7 | x |
|  | NCHA00086 | 2.61e4 | 9.72e7 | 3.93e8 | x |
|  | NCHA000143 | N | 7.50e6 | 1.79e6 | x |
|  | NCHA000150 | 1.01e4 | 1.35e8 | 1.48e8 | x |
|  | NCHA000178 | 1.33e6 | 4.88e9 | 8.63e8 | 2.26e4 |
|  | NCHA000180 | 1.58e5 | 3.29e8 | 6.94e8 | x |
|  | NCHA000181 | 6.77e4 | 8.33e7 | 7.00e7 | 2.08e4 |

x : samples not available. N indicates that the qRT-PCR has been done but the cycle threshold was too high (above 45).

**Supplementary Table 3.** Anti-PUUV antibody responses of bank voles to PUUV infection through time.

| Region | Individual number | OD<sub>450nm</sub> |  |  |  |  |  |  |  |  |
| --- | --- | --- | --- | --- | --- | --- | --- | --- | --- | --- |
|  |  | 0 dpi | 8 dpi | 14 dpi | 20 dpi | 28 dpi | 36 dpi | 43 dpi | 49 dpi | 55 dpi |
| Ain<br>non-endemic | 15.129_63 | 0.0000 | 0.0050 | 0.0700 | 0.0890 | **0.1270** | **0.2440** | **0.258** | **0.24** | **0.264** |
|  | 15.129_67 | 0.0010 | 0.0025 | 0.0000 | **0.1090** | **0.1640** | **0.1510** | **0.155** | **0.144** | **0.118** |
|  | 15.129_89 | 0.0000 | 0.0040 | 0.0380 | 0.0320 | 0.0750 | **0.1270** | **0.146** | **0.137** | **0.135** |
|  | 15.129_92 | 0.0020 | 0.0210 | 0.0810 | **0.1430** | **0.1550** | **0.1650** | **0.217** | **0.172** | **0.196** |
|  | 15.129_96 | 0.0030 | 0.0040 | 0.0050 | 0.0380 | 0.0700 | **0.1540** | **0.172** | **0.178** | **0.129** |
|  | 15.129_98 | 0.0020 | 0.0675 | **0.1240** | **0.1910** | **0.1840** | **0.2050** | **0.173** | **0.154** | **0.172** |
|  | 15.129_100 | 0.0030 | 0.0025 | 0.0090 | 0.0380 | 0.0850 | 0.0520 | 0.034 | 0.039 | 0.04 |
|  | 15.129_66 | 0.0010 | 0.0310 | 0.0970 | 0.0420 | 0.0100 | 0.0150 | 0.018 | 0.028 | 0.042 |
| Jura<br>endemic | 15.131_52 | 0.0000 | 0.0410 | 0.0850 | **0.1180** | **0.1120** | 0.0870 | 0.097 | 0.061 | 0.099 |
|  | 15.131_53 | 0.0060 | 0.0615 | **0.1710** | **0.2450** | **0.2460** | **0.2950** | **0.441** | **0.405** | **0.27** |
|  | 15.131_68 | 0.0050 | 0.0600 | **0.1020** | **0.1490** | **0.2550** | **0.3260** | **0.467** | **0.308** | **0.241** |
|  | 15.131_70 | 0.0000 | 0.0310 | **0.1150** | **0.1330** | **0.2580** | **0.1890** | **0.285** | **0.25** | **0.206** |
|  | 15.131_71 | 0.0070 | 0.0015 | **0.1310** | **0.1430** | **0.2550** | **0.2630** | **0.296** | **0.222** | **0.257** |
|  | 15.131_72 | 0.0060 | 0.0925 | **0.1220** | **0.1830** | **0.1950** | **0.1820** | **0.235** | **0.213** | **0.198** |
|  | 15.131_73 | 0.0000 | 0.0710 | 0.0950 | **0.1060** | **0.1680** | **0.1500** | **0.216** | **0.098** | **0.143** |
|  | 15.131_74 | 0.0000 | 0.0665 | **0.1220** | **0.1430** | **0.1640** | **0.1550** | **0.173** | **0.175** | **0.202** |

The amount of anti-PUUV IgG is expressed using the optical density measured at 450 nm (OD<sub>450nm</sub>). A sample is considered as seropositive when the OD<sub>450nm</sub> exceeds 0.1 (values are then indicated in bold).
The three animals that died at 28 dpi are not included in the Table.

**Supplementary Table 4.** Detection of PUUV RNA in excretas collected at 8 and 20 days post-infection using nested RT-PCR.

| Region | Individual number | Serological status | Sera |  | Feces |  | Saliva |  |
| --- | --- | --- | --- | --- | --- | --- | --- | --- |
|  |  |  | 8dpi | 20dpi | 8dpi | 20dpi | 8dpi | 20dpi |
| Ain<br>non-endemic | 15.129_63 | + | POS | N | N | N | N | N |
|  | 15.129_67 | + | N | N | N | N | N | N |
|  | 15.129_89 | + | N | N | N | N | N | N |
|  | 15.129_92 | + | N | N | N | N | N | N |
|  | 15.129_96 | + | N | POS | N | N | N | N |
|  | 15.129_98 | + | N | N | N | N | N | N |
|  | 15.129_100 | - | N | N | N | N | N | N |
|  | 15.129_66 | - | POS | N | N | N | N | N |
| Jura endemic | 15.131_52 | + | POS | N | N | N | N | N |
|  | 15.131_53 | + | N | N | N | N | N | N |
|  | 15.131_68 | + | POS | N | N | N | N | N |
|  | 15.131_70 | + | N | N | N | N | N | N |
|  | 15.131_71 | + | N | N | N | N | N | N |
|  | 15.131_72 | + | POS | N | N | N | N | N |
|  | 15.131_73 | + | POS | N | N | N | N | N |
|  | 15.131_74 | + | N | N | N | N | N | N |

- indicates bank voles that did not seroconvert during the experiment; + indicates bank voles that seroconverted during the experiment. POS indicates that the nested RT-PCR enabled to amplify PUUV RNA.

**Supplementary Table 5.** Detection of PUUV RNA in organs and excretas collected 55 days after experimental infections using quantitative RT-PCR.

| Region | Individual number | Serological status | Sera | Lung | Liver | Kidney | Saliva | Urine | Feces |
| --- | --- | --- | --- | --- | --- | --- | --- | --- | --- |
| Ain<br>non-endemic | 15.129_63 | + | N | 4.07e7 | 3.07e6 | 1.76e6 | N | N | N |
|  | 15.129_67 | + | N | 4.34e5 | N | N | N | N | N |
|  | 15.129_89 | + | N | 5.03e6 | 1.11e6 | 1.48e6 | N | N | N |
|  | 15.129_92 | + | N | N | N | N | N | N | N |
|  | 15.129_96 | + | N | 3.32e6 | 1.69e6 | 1.71e7 | N | N | 1.64e7 |
|  | 15.129_98 | + | N | 4.89e6 | N | 8.59e5 | N | N | N |
|  | 15.129_100 | - | N | N | N | N | N | N | N |
|  | 15.129_66 | - | N | 7.65e6 | N | 9.58e6 | N | N | N |
| Jura endemic | 15.131_52 | + | N | 1.03e7 | N | N | N | N | N |
|  | 15.131_53 | + | N | 6.35e6 | N | 2.00e6 | N | N | 3.55e6 |
|  | 15.131_68 | + | N | 3.54e6 | N | 5.30e5 | N | N | N |
|  | 15.131_70 | + | N | 4.30e6 | N | N | N | N | N |
|  | 15.131_71 | + | N | 3.17e7 | 1.32e6 | 6.50e5 | N | N | N |
|  | 15.131_72 | + | N | 8.90e5 | N | 5.19e5 | N | N | N |
|  | 15.131_73 | + | N | N | N | 4.35e5 | N | N | N |
|  | 15.131_74 | + | N | 5.74e5 | N | N | N | N | N |

- indicates bank voles that did not seroconvert during the experiment; + indicates bank voles that seroconverted during the experiment. Viral RNA is expressed in RNA copies. mg<sup>-1</sup>. N indicates that the qRT-PCR has been done but the cycle threshold was too high (above 45).

## References

Aars, J., Dallas, J.F., Piertney, S.B., Marshall, F., Gow, J.L., Telfer, S., Lambin, X., 2006. Widespread gene flow and high genetic variability in populations of water voles *Arvicola terrestris* in patchy habitats. Mol. Ecol. 15, 1455–1466.

Behnke, J.M., Barnard, C.J., Bajer, A., Bray, D., Dinmore, J., Frake, K., Osmond, J., Race, T., Sinski, E., 2001. Variation in the helminth community structure in bank voles (*Clethrionomys glareolus*) from three comparable localities in the Mazury Lake District region of Poland. Parasitol. 123, 401–414.

Beldomenico, P.M., Telfer, S., Gebert, S., Lukomski, L., Bennett, M., Begon, M., 2008. Poor condition and infection: a vicious circle in natural populations. P. R. Soc. Lond. B 275, 1753–1759.

Benjamini, Y., Hochberg, Y., 1995. Controlling the false discovery rate: a practical and powerful approach to multiple testing. J. R. Stat. Soc. B 57, 289–300.

Bernshtein, A.D., Apekina, N.S., Mikhailova, T.V., Myasnikov, Y.A., Khlyap, L.A., Korotkov, Y.S., Gavrilovskaya, I.N., 1999. Dynamics of Puumala hantavirus infection in naturally infected bank voles (*Clethrinomys glareolus*). Arch. Virol. 144, 2415–2428.

Berthier, K., Charbonnel, N., Galan, M., Chaval, Y., Cosson, J.F., 2006. Migration and maintenance of genetic variability in cyclic vole populations : a spatio-temporal study. Mol. Ecol. 15, 2665–2676.

Berthier, K., Galan, M., Foltete, J.C., Charbonnel, N., Cosson, J.F., 2005. Genetic structure of the cyclic fossorial water vole (*Arvicola terrestris*): landscape and demographic influences. Mol. Ecol. 14, 2861–2871.

Bowie, A.G., Haga, I.R., 2005. The role of Toll-like receptors in the host response to viruses. Mol. Immunol. 42, 859–867.

Bradley, J.R., 2008. TNF-mediated inflammatory disease. J. Pathol. 214, 149-160.

Brummer-Korvenkotio, M., Henttonen, H., Vaheri, A., 1982. Hemorrhagic fever with renal syndrome in Finland: Ecology and virology of nephropathia epidemica. Scand. Journal of Infect. Dis. 36, 88–89.

Bryja, J., Charbonnel, N., Berthier, K., Galan, M., Cosson, J.F., 2007. Density-related changes in selection pattern for major histocompatibility complex genes in fluctuating populations of voles. Mol. Ecol. 16, 5084–5097.

Castel, G., Couteaudier, M., Sauvage, F., Pons, J.B., Murri, S., Plyusnina, A., Pontier, D., Cosson, J.F., Plyusnin, A., Marianneau, P., Tordo, N., 2015. Complete genome and phylogeny of Puumala hantavirus isolates circulating in France. Viruses 7, 5476–5488.

Charbonnel, N., Pagès, M., Sironen, T., Henttonen, H., Vapalahti, O., Mustonen, J., Vaheri, A., 2014. Immunogenetic factors affecting susceptibility of humans and rodents to hantaviruses and the clinical course of hantaviral disease in humans. Viruses 6, 2214–2241.

Daszak, P., Cunningham, A.A., Hyatt, A.D., 2000. Emerging infectious diseases of wildlife - threats to biodiversity and human health. Science 287, 443–449.

Deter, J., Chaval, Y., Galan, M., Gauffre, B., Morand, S., Henttonen, H., Laakkonen, J., Voutilainen, L., Charbonnel, N., Cosson, J.F., 2008a. Kinship, dispersal and hantavirus transmission in bank and common voles. Arch. Virol. 153, 435–444.

Deter, J., Chaval, Y., Galan, M., Henttonen, H., Laakkonen, J., Voutilainen, L., Ribas Salvador, A., Bryja, J., Morand, S., Cosson, J.F., Charbonnel, N., 2008b. Association between the DQA MHC class II gene and Puumala virus infection in the specific reservoir Myodes glareolus. Inf. Genet. Evol. 8, 450–458.

Di Ciccio, T.J., Efron, B., 1996. Bootstrap confidence intervals. Stat. Sci. 11, 189–228.

Dubois, A., Castel, G., Murri, S., Pulido, C., Pons, J.B., Benoit, L., Loiseau, A., Lakhdar, L., Galan, M., Charbonnel, N., Marianneau, P., 2017a. Experimental infections of wild bank voles (*Myodes glareolus*) from nephropatia epidemica endemic and non-endemic regions revealed slight differences in Puumala virological course and immunological responses. Virus Res.

Dubois, A., Galan, M., Guivier, E., Henttonen, H., Voutilainen, L., Razzauti, M., Gauffre, B., Marianneau, P., Cosson, J.F., Charbonnel, N., 2017b. Microevolution of bank voles (*Myodes glareolus*) at neutral and immune related genes during a multi-annual complete dynamic cycle : consequences for Puumala hantavirus epidemiology. Inf. Genet. Evol. 49, 318–329.

Easterbrook, J.D., Klein, S.L., 2008. Seoul virus enhances regulatory and reduces proinflammatory responses in male Norway rats. J. Med. Virol. 80, 1308–1318.

Excoffier, L., Lischer, H.E.L., 2010. Arlequin suite ver 3.5: a new series of programs to perform population genetics analyses under Linux and Windows. Mol. Ecol. Res. 10, 564–567.

Friberg, I.M., Lowe, A., Ralli, C., Bradley, J.E., Jackson, J.A., 2011. Temporal anomalies in immunological gene expression in a time series of wild mice: signature of an epidemic? PLoS One 6.

Gascuel, F., Choisy, M., Duplantier, J.M., Debarre, F., Brouat, C., 2013. Host resistance, population structure and the long-term persistence of bubonic plague: Contributions of a modelling approach in the malagasy focus. Plos Comput. Biol. 9, e1003039.

Gavrilovskaya, I.N., Chumakov, M.P., Apekina, N.S., Ryltseva, E.V., Martiyanova, L.I., Gorbachkova, E.A., Bernshtein, A.D., Zakharova, M.A., Boiko, V.A., 1983. Adaptation to laboratory and wild animals of the haemorrhagic fever with renal syndrome virus present in the foci of European U.S.S.R. Arch. Virol. 77, 87–90.

Guivier, E., Galan, M., Chaval, Y., Xuereb, A., Ribas Salvador, A., Poulle, M.L., Charbonnel, N., Cosson, J.F., 2011. Landscape genetics highlights the role of bank vole metapopulation dynamics in the epidemiology of Puumala hantavirus. Mol. Ecol. 20, 3569–3583.

Guivier, E., Galan, M., Henttonen, H., Cosson, J.F., Charbonnel, N., 2014. Landscape features and helminth co-infection shape bank vole immunoheterogeneity, with consequences for Puumala virus epidemiology. Heredity 112, 274–281.

Guivier, E., Galan, M., Ribas Salvador, A., Xuéreb, A., Chaval, Y., Olsson, G., Essbauer, S., Henttonen, H., Voutilainen, L., Cosson, J.F., Charbonnel, N., 2010. Tnf-a expression and promoter sequences reflect the balance of tolerance/resistance to Puumala virus infection in European bank vole populations. Inf. Genet. Evol. 10, 1208–1217.

Hardestam, J., Karlsson, M., Falk, K.I., Olsson, G., Klingström, J., Lundkvist, A., 2008. Puumala hantavirus excretion kinetics in bank voles (*Myodes glareolus*). Emerg. Infect. Dis. 14, 1209–1215.

Hayward, A.D., Nussey, D.H., Wilson, A.J., Berenos, C., Pilkington, J.G., Watt, K.A., Pemberton, J.M., Graham, A.L., 2014. Natural selection on individual variation in tolerance of gastrointestinal nematode infection. Plos Biology 12, e1001917.

Heil, F., Hemmi, H., Hochrein, H., Ampenberger, F., Kirschning, C., Akira, S., Lipford, G., Wagner, H., Bauer, S., 2004. Species-specific recognition of single-stranded RNA via toll-like receptor 7 and 8. Science 303, 1526–1529.

Henttonen, H., Buchy, P., Suputtamongkol, Y., Jittapalapong, S., Herbreteau, V., Laakkonen, J., Chaval, Y., Galan, M., Dobigny, G., Charbonnel, N., Michaux, J., Cosson, J.F., Morand, S., Hugot, J.P., 2008. Recent discoveries of new hantaviruses widen their range and question their origins, in: Sparagano, O.A.E., Maillard, J.C., Figueroa, J.V. (Eds.), Animal biodiversity and emerging diseases: prediction and prevention. Blackwell Publishing, Oxford, pp. 84–89.

Heyman, P., Ceianu, C.S., Christova, I., Tordo, N., Beersma, M., Joao Alves, M., Lundkvist, A., Hukic, M., Papa, A., Tenorio, A., Zelena, H., Essbauer, S., Visontai, I., Golovljova, I., Connell, J., Nicoletti, L., Van Esbroeck, M., Gjeruldsen Dudman, S., Aberle, S.W., Avsic-Zupanc, T., Korukluoglu, G., Nowakowska, A., Klempa, B., Ulrich, R.G., Bino, S., Engler, O., Opp, M., Vaheri, A., 2011. A five-year perspective on the situation of haemorrhagic fever with renal syndrome and status of the hantavirus reservoirs in Europe, 2005-2010. Eurosurveillance 16.

Heyman, P., Thoma, B.R., Marie, J.L., Cochez, C., Essbauer, S.S., 2012. In search for factors that drive hantavirus epidemics. Front. Physiol. 3, 237.

Hirsch, I., Caux, C., Hasan, U., Bendriss-Vermare, N., Olive, D., 2010. Impaired Toll-like receptor 7 and 9 signaling : from chronic viral infections to cancer. Trends Immunol. 31, 391-397.

Jackson, J.A., Begon, M., Birtles, R., Paterson, S., Friberg, I.M., Hall, A., Ralli, C., Turner, A., Zawadzka, M., Bradley, J.E., 2011. The analysis of immunological profiles in wild animals: a case study on immunodynamics in the field vole, *Microtus agrestis*. Mol. Ecol. 20, 893-909.

Jackson, J.A., Hall, A.J., Friberg, I.M., Ralli, C., Lowe, A., Zawadzka, M., Turner, A.K., Stewart, A., Birtles, R.J., Paterson, S., Bradley, J.E., Begon, M., 2014. An immunological marker of tolerance to infection in wild rodents. Plos Biol. 12, e1001901.

Jameson, L.H., Logue, C.H., Atkinson, B., Baker, B., Galbraith, S.E., Carroll, M.W., Brooks, T., Hewson, R., 2013. The continued emergence of hantaviruses: isolation of a Seoul virus implicated in human disease, United Kingdom, October 2012. Eurosurveillance 18.

Jin, H.K., Yoshimatsu, K., Takada, A., Ogino, M., Asano, A., Arikawa, J., Watanabe, T., 2001. Mouse Mx2 protein inhibits hantavirus but not influenza virus replication. Arch. Virol. 146, 41–49.

Jombart, T., Devillard, S., Balloux, F., 2010. Discriminant analysis of principal components: a new method for the analysis of genetically structured populations. BMC Genet. 11, 94.

Kallio, E.R., Klingström, J., Gustafsson, E., Manni, T., Vaheri, A., Henttonen, H., Vapalahti, O., Lundkvist, A., 2006. Prolonged survival of Puumala hantavirus outside the host: evidence for indirect transmission via the environment. J. Gen. Virol. 87, 2127–2134.

Kanerva, M., Melen, K., Vaheri, A., Julkunen, I., 1996. Inhibition of Puumala and Tula hantaviruses in vero cells by MxA protein. Virol. 224, 55–62.

King, D.A., Peckham, C., Waage, J.K., Brownlie, J., Woolhouse, M.E., 2006. Epidemiology. Infectious diseases: preparing for the future. Science 313, 1392–1393.

Klein, S.L., Flanagan, K.L., 2016. Sex differences in immune responses. Nat. Rev. Immunol. 16, 626–638.

Klingstrom, J., Heyman, P., Escutenaire, S., Sjolander, K.B., De Jaegere, F., Henttonen, H., Lundkvist, A., 2002. Rodent host specificity of European hantaviruses: evidence of Puumala virus interspecific spillover. J. Med. Virol. 68, 581–588.

Korpela, H., Lähdevirta, J., 1978. The role of small rodents and patterns of living in the epidemiology of nephropathia epidemica. Scand. J. Infect. Dis. 10, 303–305.

Korva, M., Duh, D., Saksida, A., Trilar, T., Avsic-Zupanc, T., 2009. The hantaviral load in tissues of naturally infected rodents. Microb. Infect. 11, 344–351.

Kruger, D.H., Ulrich, R., Lundkvist, Å., 2001. Hantavirus infections and their prevention. Microb. Infect. 3, 1129–1144.

Lee, K.A., Klasing, K.C., 2004. A role for immunology in invasion biology. Trends Ecol. Evol. 19, 523–529.

Li, Y.L., Youssoufian, H., 1997. MxA overexpression reveals a common genetic link in four Fanconi anemia complementation groups. J. Clin. Invest. 100, 2873–2880.

Lloyd-Smith, J.O., Cross, P.C., Briggs, C.J., Daugherty, M., Getz, W.M., Latto, J., Sanchez, M.S., Smith, A.B., Swei, A., 2005. Should we expect population thresholds for wildlife disease? Trends Ecol. Evol. 20, 511–519.

Loxton, K.C., Lawton, C., Stafford, P., Holland, C.V., 2016. Reduced helminth parasitism in the introduced bank vole (*Myodes glareolus*): More parasites lost than gained. Int. J. Parasitol. 5, 175–183.

Lund, J.M., Alexopoulou, L., Sato, A., Karow, M., Adams, N.C., Gale, N.W., Iwasaki, A., Flavell, R.A., 2004. Recognition of single-stranded RNA viruses by Toll-like receptor 7. P. Nat. Acad. Sci. USA 101, 5598–5603.

Mace, G., Feyeux, C., Mollard, N., Chantegret, C., Audia, S., Rebibou, J.M., Spagnolo, G., Bour, J.B., Denoyel, G.A., Sagot, P., Reynes, J.M., 2013. Severe Seoul hantavirus infection in a pregnant woman, France, October 2012. Eurosurveillance 18, 14–17.

Morse, S.S., Schluederberg, A., 1990. Emerging viruses: the evolution of viruses and viral diseases. J. Infect. Dis. 162, 1–7.

Olsson, G.E., White, N., Ahlm, C., Elgh, F., Verlemyr, A.C., Juto, P., Palo, R.T., 2002. Demographic factors associated with hantavirus infection in bank voles (*Clethrionomys glareolus*). Emerg. Infect. Dis. 8, 924–929.

Pfaffl, M.W., 2001. A new mathematical model for relative quantification in real-time RT-PCR. Nucl. Ac. Res. 29.

Plath, K., Mlynarczyk-Evans, S., Nusinow, D.A., Panning, B., 2002. Xist RNA and the mechanism of X chromosome inactivation. Annu. Rev. Genet. 36, 233–278.

Raberg, L., Graham, A.L., Read, A.F., 2009. Decomposing health: tolerance and resistance to parasites in animals. Philos. Trans. R. Soc. B-Biol. Sci. 364, 37–49.

Raymond, M., Rousset, F., 1995. Genepop version 3: population genetics software for exact tests and ecumenicism. J. Heredity 86, 248–249.

Razzauti-Feliu, M., Galan, M., Bernard, M., Maman, S., Klopp, C., Charbonnel, N., Vayssier-Taussat, M., Eloit, M., Cosson, J.F., 2015. Comparison of next-generation sequencing approaches surveying bacterial pathogens in wildlife. Plos Negl. Trop. Dis. 9, e0003929.

Reynes, J.M., Dutrop, C.M., Carli, D., Levast, M., Fontaine, N., Denoyel, G.A., Philit, J.B., 2015. Puumala hantavirus infection in Isere: Geographic extension of this zoonosis in France. Med. Malad. Infect. 45, 177–180.

Ribas Salvador, A., Guivier, E., Xuereb, A., Chaval, Y., Cadet, P., Poulle, M.L., Sironen, T., Voutilainen, L., Henttonen, H., Cosson, J.F., Charbonnel, N., 2011. Concomitant influence of helminth infection and landscape on the distribution of Puumala hantavirus in its reservoir, *Myodes glareolus*. BMC Microbiol. 11.

Rikalainen, K., Grapputo, A., Knott, E., Koskela, E., Mappes, T., 2008. A large panel of novel microsatellite markers for the bank vole (*Myodes glareolus*). Mol. Ecol. Res. 8, 1164–1168.

Rohfritsch, A., Guivier, E., Galan, M., Chaval, Y., Charbonnel, N., 2013. Apport de l’immunogénétique à la compréhension des interactions entre le campagnol roussâtre *Myodes glareolus* et l’hantavirus Puumala. Bull. Acad. Vet. France 166, 165–176.

Ruijter, J.M., Ramakers, C., Hoogaars, W.M.H., Karlen, Y., Bakker, O., van den Hoff, M.J.B., Moorman, A.F.M., 2009. Amplification efficiency: linking baseline and bias in the analysis of quantitative PCR data. Nucl. Ac. Res. 37.

Schneider, D.S., Ayres, J.S., 2008. Two ways to survive infection: what resistance and tolerance can teach us about treating infectious diseases. Nat. Rev. Immunol. 8, 889–895.

Schonrich, G., Rang, A., Lutteke, N., Raftery, M.J., Charbonnel, N., Ulrich, R.G., 2008. Hantavirus-induced immunity in rodent reservoirs and humans. Immunol. Rev. 225, 163–189.

Schountz, T., Acuna-Retamar, M., Feinstein, S., Prescott, J., Torres-Perez, F., Podell, B., Peters, S., Ye, C., Black, W.C., Hjelle, B., 2012. Kinetics of immune responses in deer mice experimentally infected with Sin Nombre Virus. J. Virol. 86, 10015–10027.

Schountz, T., Prescott, J., 2014. Hantavirus immunology of rodent reservoirs: current status and future directions. Viruses 6, 1317–1335.

Sommaruga, A., 1997. Geology of the Central Jura and the Molasse Basin, Institut de Géologie. Université de Neuchâtel, Neuchâtel, Suisse, p. 195.

Temonen, M., Mustonen, J., Helin, H., Pasternack, A., Vaheri, A., Holthofer, H., 1996. Cytokines, adhesion molecules, and cellular infiltration in nephropathia epidemica kidneys: An immunohistochemical study. Clin. Immunol. Immunopathol. 78, 47–55.

Tersago, K., Crespin, L., Verhagen, R., Leirs, H., 2012. Impact of Puumala virus infection on maturation and survival in bank voles: A capture-mark-recapture analysis. J. Wildl. Dis. 48, 148–156.

Tollenaere, C., Jacquet, S., Ivanova, S., Loiseau, A., Duplantier, J.M., Streiff, R., Brouat, C., 2013. Beyond an AFLP genome scan towards the identification of immune genes involved in plague resistance in Rattus rattus from Madagascar. Mol. Ecol. 22, 354–367.

Vaheri, A., Strandin, T., Hepojoki, J., Sironen, T., Henttonen, H., Mäkelä, S., Mustonen, J., 2013. Uncovering the mysteries of hantavirus infections. Nat. Rev. Microbiol. 11, 539–550.

Vander Wal, E., Garant, D., Calmé, S., Chapman, C.A., Festa-Bianchet, M., Millien, V., Rioux-Paquette, S., Pelletier, F., 2014. Applying evolutionary concepts to wildlife disease ecology and management. Evol. Applic. 7, 856–868.

Vapalahti, O., Mustonen, J., Lundkvist, A., Henttonen, H., Plyusnin, A., Vaheri, A., 2003. Hantavirus infections in Europe. Lancet Inf. Dis. 3, 653–661.

Voutilainen, L., Kallio, E.R., Niemimaa, J., Vapalahti, O., Henttonen, H., 2016. Temporal dynamics of Puumala hantavirus infection in cyclic populations of bank voles. Sci. Rep. 6, 21323.

Voutilainen, L., Sironen, T., Tonteri, E., Tuiskunen Back, A., Razzauti, M., Karlsson, M., Wahlstrom, M., Niemimaa, J., Henttonen, H., Lundkvist, A., 2015. Life-long shedding of Puumala hantavirus in wild bank voles (*Myodes glareolus*). J. Gen. Virol. 96, 1238–1247.

Weir, B., Cockerham, C., 1984. Estimating F-statistics for the analysis of population structure. Evolution 38, 1358–1370.

Wenzel, J., Uerlich, M., Haller, O., Bieber, T., Tueting, T., 2005. Enhanced type I interferon signaling and recruitment of chemokine receptor CXCR3-expressing lymphocytes into the skin following treatment with the TLR7-agonist imiquimod. J. Cutan. Pathol. 32, 257–262.

Wright, S., 1951. The genetical structure of populations. Ann. Eugenics 15, 323–354.

Yanagihara, R., Amyx, H.L., Gajdusek, D.C., 1985. Experimental infection with Puumala virus, the etiologic agent of nephropathia epidemica, in bank voles (*Clethrionomys glareolus*). J. Virol. 55, 34–38.

Zeimes, C.B., Quoilin, S., Henttonen, H., Lyytikaïnen, O., Vapalahti, O., Reynes, J.M., Reusken, C., Swart, A.N., Vainio, K., Hjertqvist, M., Vanwambeke, S.O., 2015. Landscape and regional environmental analysis of the spatial distribution of hantavirus human cases in Europe. Front. Publ. Health 3, 1–10.

